# Distinct roles of IDR and CCD domains control PABPN1 aggregation and enable therapeutic rescue

**DOI:** 10.64898/2026.06.12.731084

**Authors:** Milad Shademan, Iolanda Vendrell, Alberto Malerba, Sogol Khoshbash, Sarah Flannery, James March, Geertje van der Horst, Silvère M. van der Maarel, Benedikt M. Kessler, Vered Raz

## Abstract

PABPN1 is a multifunctional protein whose expression is tightly regulated to maintain cellular homeostasis. PABPN1 dysregulation contributes to both common acquired diseases, such as bladder cancer, and rare inherited disorders, including oculopharyngeal muscular dystrophy (OPMD). In OPMD, PABPN1 forms insoluble aggregates that reduce its functional levels, leading to genome-wide shifts in alternative polyadenylation (APA) at 3′-UTRs and disruption of mRNA metabolism, including nuclear export and translation. OPMD is caused by a short alanine expansion at the N-terminus of PABPN1 within an intrinsically disordered region (IDR) followed by a coiled-coil domain (CCD). How these domains contribute to PABPN1 function and aggregation remains unclear.

Here, we show that the N-terminal IDR promotes aggregation and modulates protein-protein interactions with longer IDRs suppressing interaction between PABPN1 and its binding partners. We further demonstrate that the CCD has dual functions: its N-terminal domain dictates PABPN1 stability, whereas its C-terminal domain enhances the PABPN1 interactome. We identified a naturally occurring variant lacking exon 1 encoding the IDR and the CCD N-terminal domain (named trPAB). That variant forms a stable, non-aggregating protein isoform. Interactome analysis and structural modeling indicate improved molecular function by restoring PABPN1 activity, APA profiles, and cellular phenotypes in OPMD and bladder cancer models. Importantly, delivery of trPAB via Adeno-associated viral vector in an OPMD mouse model improves muscle histopathology, supporting its potential as a therapeutic strategy.

**A mechanistic basis of a novel gene therapy approach for OPMD:** 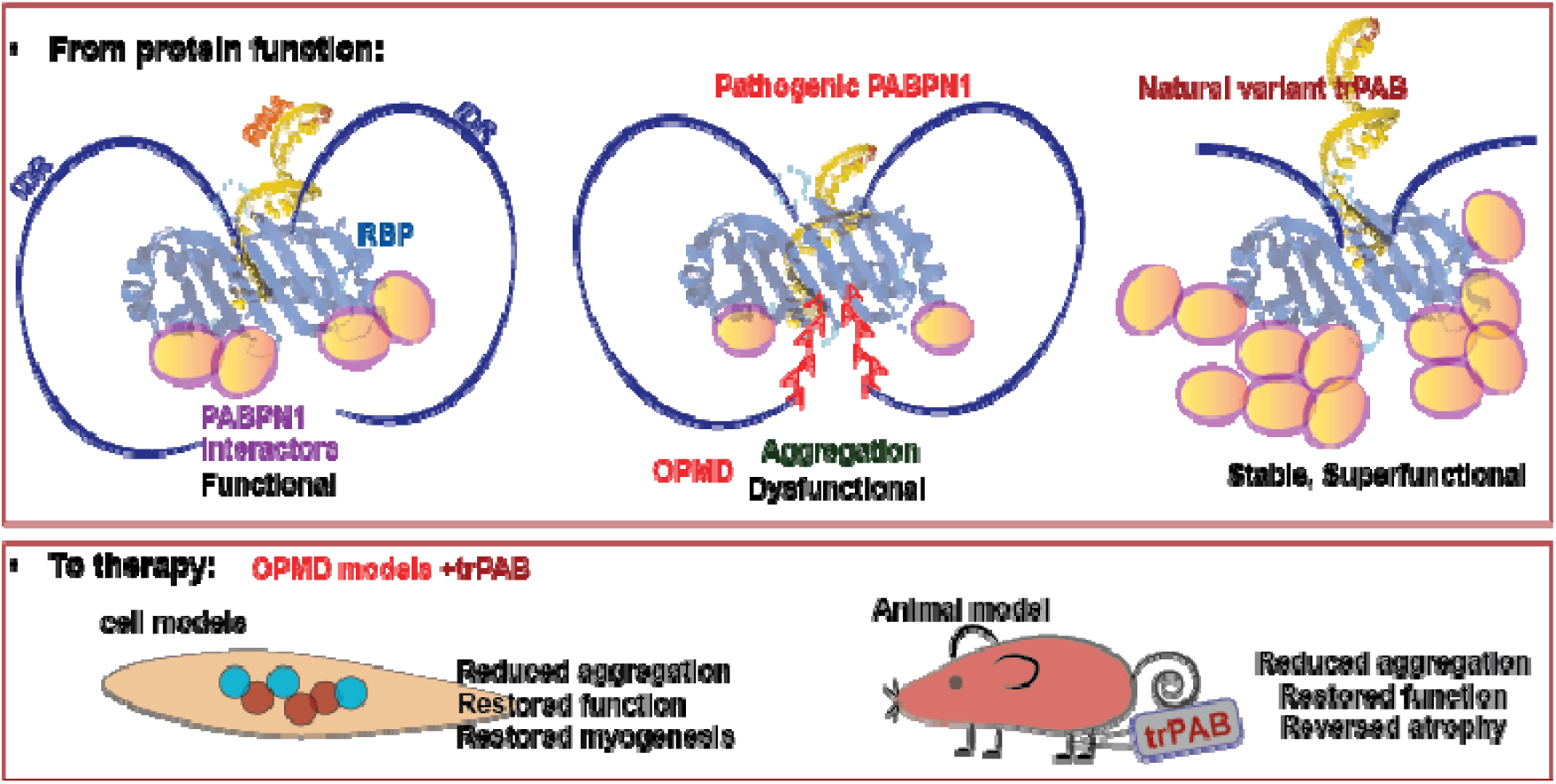

Top panel: A schematic presentation of three PABPN1 variant functional states. Left: the normal full-length protein interacts with key RNA binding protein (RBP) partners supporting normal cellular function. Middle: A pathogenic full-length alanine-expanded PABPN1 leads to protein aggregation, resulting in limited interactors, dysfunctional complexes and impaired cellular processes. Right: a truncated natural variant (trPAB), not associated with OPMD pathology, shows enhanced stability and reduced aggregation, suggesting a potential protective or “super functional” profile.

The bottom panel translates these mechanistic insights into therapeutic strategy. In cell models, expression of trPAB is associated with reduced protein aggregation, restoration of normal cellular function, and improved myogenesis. These effects are recapitulated in a relevant animal model of OPMD, where trPAB treatment leads to decreased aggregation, functional recovery, and reversal of muscle atrophy.

Our study supports the rationale for leveraging a stabilized PABPN1 variant as a novel gene therapy approach to counteract aggregation-driven pathology in OPMD.

**Highlights:**

- The central coiled-coil domain (CCD) of PABPN1 has two opposing functions: its N-terminal region stabilizes the protein, while its C-terminal region promotes stability but hampers aggregation.
- At the N-terminus, an intrinsically disordered region (IDR) acts as a gatekeeper for protein interactions. The alanine tract, including its pathogenic expansion, restricts this interactome, limiting PABPN1 binding capacity.
- We identified a naturally occurring PABPN1 isoform, trPAB, which lacks exon 1, including the IDR and the N-terminal portion of the CCD. trPAB forms a stable and functional protein and exerts beneficial effects in muscle cells.
- Functionally, trPAB rescues key cellular phenotypes in bladder cancer cells and reverses nuclear aggregation and muscle atrophy in a mouse model of OPMD.

## Introduction

RNA-binding proteins (RBPs) are key regulators of mRNA fate, controlling processing, export, stability, and translation. Disruption of RBP function is increasingly recognized as a unifying mechanism across diverse diseases, particularly in neuromuscular disorders ^1^. A defining feature of many RBPs is their intrinsic tendency to aggregate, which can lead to dysfunction ^2^. All RBPs contain an RNA-binding motif or domain (RBD); the canonical RBD adopts a structure composed of a four-stranded β-sheet packed against two α-helices ^3^. RBP aggregation is often influenced by mutations, including point mutations and trinucleotide expansions, which promote misfolding and impair function ^1,4,5^. These mutations frequently occur in intrinsically disordered regions (IDRs), which lack a defined structure and often serve as flexible linkers between structured domains ^6^. Although aggregation is widely associated with pathology, the structural features that link RBP organization to function and disease remain poorly understood.

Poly(A)-binding protein nuclear 1 (PABPN1) exemplifies this paradigm. PABPN1 is an essential and evolutionarily conserved regulator of mRNA processing and nuclear export ^7^. A short alanine expansion (+1 to +8) within the normal 10-alanine tract at the N-terminus is sufficient to cause oculopharyngeal muscular dystrophy (OPMD), a late-onset myopathy with striking tissue specificity ^8^. Despite its ubiquitous expression, it remains unclear why PABPN1 dysfunction selectively affects skeletal muscle.

A disease mechanism in which PABPN1 aggregation drives functional depletion, reducing its expression levels has been proposed by us and others ^7,9^, and emerging evidence supports this model ^10,11^. Reduced PABPN1 expression has also been linked to different cancers, including bladder cancer ^12,13^, suggesting that PABPN1 dosage is tightly coupled to cellular homeostasis. Although predominantly nuclear, both PABPN1 loss and aggregation impair cytosolic processes such as translation and mitochondrial activity ^14–18^, pointing to an underappreciated role in coordinating cytoplasmic mRNA function.

Structurally, PABPN1 consists of a C-terminal RNA-binding domain and a large IDR at N-terminal side that includes a short polyalanine tract followed a stable coiled-coil domain (CCD) recognized by an alpha helix topology (Fig. 1A) The CCD has been implicated in aggregation ^19–21^. A short expansion of the alanine tract is pathogenic and enhances aggregation kinetics ^22^. Structural predictions of the N-terminus in PABPN1 suggest that the alanine expanded tract may stabilize intramolecular interactions with the CCD ^22^. However, how the N-terminal IDR affect PABPN1 function is unknown.

**Figure 1.**
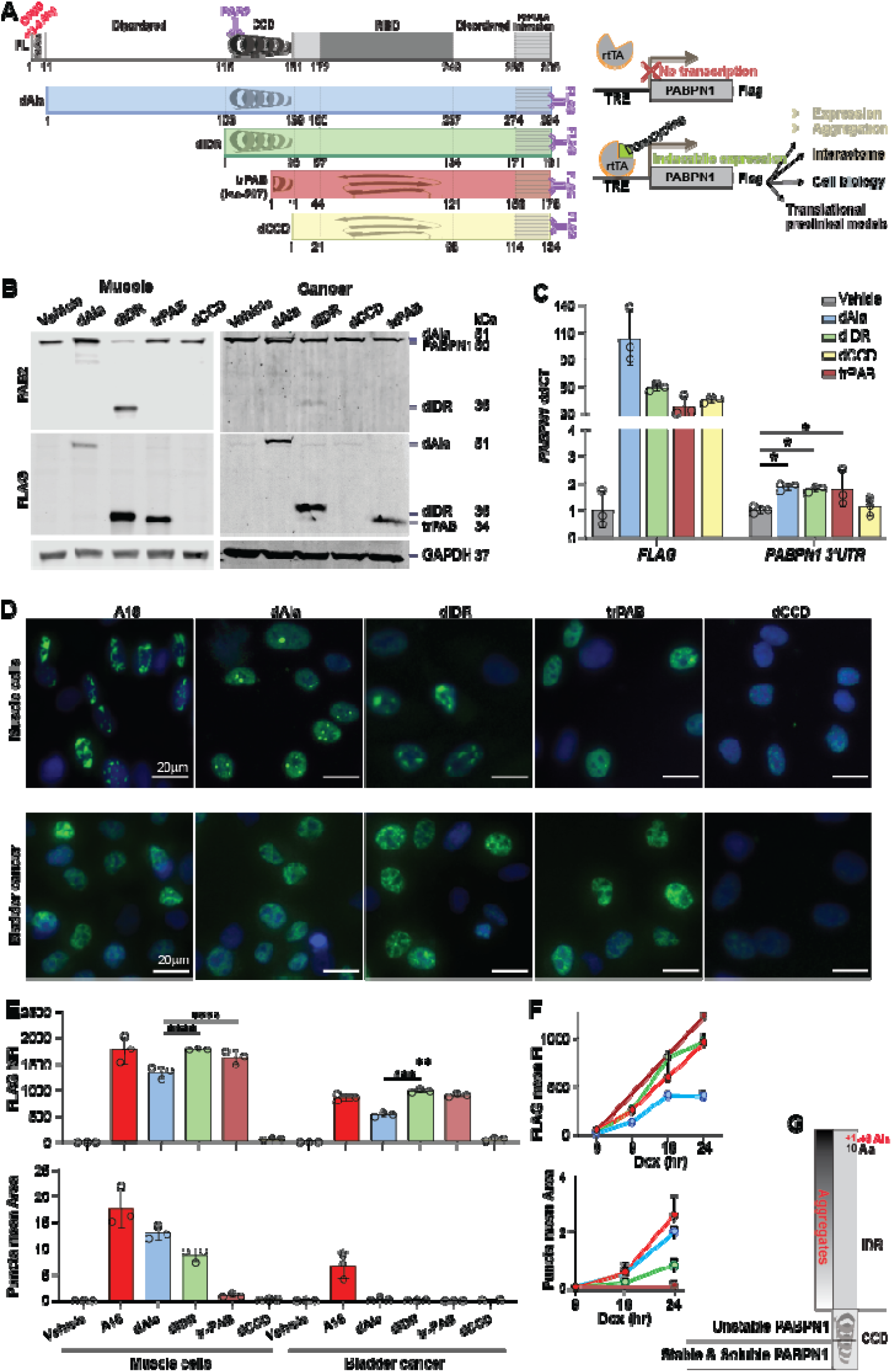
Structural characterization of the N-terminal domains of PABPN1. A. Left: Schematic representation of full-length PABPN1 (grey) and the pathogenic alanine expanded (A16) variant (red), as well as the deletions and the natural truncated variants (left). The alanine tract deletion (dAla) is shown in blue, the internal disordered region deletion (dIDR) in green, and the coiled-coil domain deletion (dCCD) in yellow. The natural isoform Iso-207, (trPAB) lacking exon 1, is highlighted in dark red. This color scheme is consistent across all panels. All variants were fused to FLAG at the C-terminus. The full length PABPN1, dAla and dIDR are also detected with the PAB2 antibody binding to the coil-coiled domain (purple). The key functional domains and predicted secondary structures are the coiled-coil domain (CCD; α-helix) and the RNA-binding domain (RBD; β-sheets and α-helix). The IDR is in light grey. Amino acid positions are provided for each domain. At the C-terminus PABPN1 binds to PAPOLA, poly(A) polymerase alpha. Right: Schematic overview of our study design: stable cells were made for each variant in muscle and cancer cells, expression is under an inducible tetracycline promoter system. The constructs were used to assess protein expression, aggregation, genome-wide interactions, cellular localization, and translational relevance in disease models. B. Representative western blots following 24 hrs Dox treatment in human muscle and bladder cancer cell lines. FLAG antibody detected induced transgene expression, while PABPN1 antibody (PAB2) detected endogenous PABPN1 as well as dAla and dIDR variants. GAPDH serves as a loading control. Molecular weights (kDa) are indicated. C. Bar chart of PABPN1 mRNA expression levels 24 hours after Dox treatment in bladder cancer cells. Transgene mRNA was detected using FLAG-specific primers, and the endogenous PABPN1 with primers targeting the 31-UTR. Values are normalized to vehicle-treated cells. Data represent mean ± SD from three biological replicates. D. Representative immunofluorescence images of FLAG staining (green) 24 hours after Dox treatment in muscle and bladder cancer cell lines. Nuclei counterstain is in blue. Scale bar: 20 μm. E. Quantification of FLAG signal 24 hours after Dox treatment in muscle and bladder cells. Upper panel shows FLAG mean nuclear fluorescence intensity (FI); Lower panel shows the mean puncta area per cell. Each dot represents the average of approximately 1,500 cells per biological replicate. F. Kinetics of FLAG fluorescence intensity and puncta formation in muscle cell lines over 24 hours of Dox treatment. G. A schematic summary illustrating the contribution of N-terminal structural regions of PABPN1 to protein accumulation and aggregation. Data are shown as mean ± SD from three biological replicates. Statistical significance was assessed using one-way ANOVA with a post-hoc Tukey test. *P* < 0.05 (*)*, < 0.01 (**), < 0.005 (***),* < 0.0001 (****).

Here, we test the hypothesis that the N-terminal region of PABPN1 acts as a structural gatekeeper of its interaction landscape. Using a series of engineered N-terminal deletions alongside a naturally occurring isoform lacking exon 1, we systematically dissect how N-terminal length and composition shape the PABPN1 interactome. We further explore the functional consequences of this regulation in both cancer and muscle contexts and assess the therapeutic potential of N-terminal truncation in cellular and animal models of OPMD.

## Results

### IDR and CCD domains determine PABPN1 stability and aggregation in a cell type–specific manner

To investigate how N-terminal architecture governs PABPN1 function, we generated a panel of deletion variants targeting the alanine tract, IDR, and CCD, alongside a naturally occurring truncated isoform (trPAB) that lacks exon 1 encoding the alanine tract, the IDR, and part of the CCD, and compared it to the pathogenic PABPN1 variant (A16) (Fig. 1A). This framework enabled systematic dissection of how discrete structural elements encode PABPN1 organization within the nucleus. All PABPN1 variants were FLAG-tagged at the C-terminus and expressed using a doxycycline-inducible system in stable cell lines, allowing controlled analysis of protein behavior across cellular contexts while minimizing potential toxicity associated with overexpression (Fig. 1A). The PAB2 antibody to the N-terminus of the CCD (Fig. 1A) detects the endogenous PABPN1, and the A16, dAla and dIDR forms, however it does not bind to dCCD and trPAB.

To probe cell type–specific effects relevant to PABPN1-associated pathology, we expressed the inducible constructs in human immortalized muscle cells and bladder cancer cells. This approach enabled comparative analysis of aggregation behaviour, cellular phenotypes, and molecular interactors across distinct cellular environments (Fig. 1A).

Across cellular contexts, all variants except the CCD deletion produced stable protein, albeit at different soluble levels, despite comparable mRNA expression across constructs (Figs. 1B and 1C). This indicates that loss of the N-terminal portion of the CCD may compromise PABPN1 protein stability. Immunofluorescence analysis showed that all variants retained nuclear localization (Fig. 1D), consistent with the mapped nuclear localization signal in the C-terminus of PABPN1^23^, and suggesting that the N-terminal region does not impair nuclear targeting, although it does exert cell type–specific effects on protein aggregation.

In muscle cells, the dAla and dIDR variants formed nuclear puncta accompanied by reduced FLAG signal, whereas puncta were not observed in bladder cancer cells, where FLAG levels remained comparable across variants (Fig. 1E). These observations suggest that PABPN1 aggregation is modulated by cell type–specific factors. In contrast to dAla and dIDR, the trPAB variant did not form puncta under any condition (Fig. 1E). Under identical induction conditions, the A16 variant formed large puncta (Figs. 1D–1E). Time-resolved analysis showed that within the first 24 hours after induction, puncta accumulation in the dAla variant was comparable to A16, whereas dIDR showed reduced puncta formation (Fig. 1F). Notably, in trPAB, high protein abundance was uncoupled from aggregation (Fig. 1F).

Collectively, these findings suggest a modular role for PABPN1 N-terminus that governs the balance between soluble and assembled nuclear states, and where the IDR length dictates aggregation (Fig. 1G). These results support a model in which N-terminal elements actively tune higher-order assembly, suggesting a role in PABPN1 function. In support, Alphafold modeling of PABPN1 folding across its variants suggested that the pathogenic PABPN1 with +3 or +6 alanine expansions fold onto the CCD and the RBD, and a shorter IDR exposed the RBD from intramolecular folding (Fig. S1), further supporting functional consequences of the IDR.

### The PABPN1 interactome is dictated by IDR length and exhibits cell type specificity

We next investigated the interactomes of PABPN1 variants in a cell type–specific manner. The trPAB interactome was assessed in both muscle and bladder cancer cells, while in muscle cells dAla, dIDR, and trPAB variants, showing pronounced differences in protein accumulation and aggregation, were analysed. The A16 variant was excluded due to its high insolubility following 24 hours of doxycycline (Dox) induction. All stable cell lines were induced with Dox for 24 hours; a time point that minimized PABPN1 insolubility and aggregation. Soluble PABPN1 variants were affinity-purified using FLAG-trap and analyzed by mass spectrometry (Fig. 2A). A principal component analysis of dataset that preprocessing of all 18 samples, muscle and cancer together, revealed that the dominant source of variance (PC1) was the cell type (Fig. 2B). Therefore, we next applied preprocessing separate for cell-type (muscle n=12 and cancer n=6) and used this dataset for all downstream analyses. In muscle cells, samples segregated along PC1, with the largest variation observed between Dox- and vehicle-treated conditions (Fig. S2). PABPN1 was the most enriched protein across all conditions, while only a few proteins were detected in vehicle controls, indicating high specificity of the FLAG-trap.

**Figure 2.**
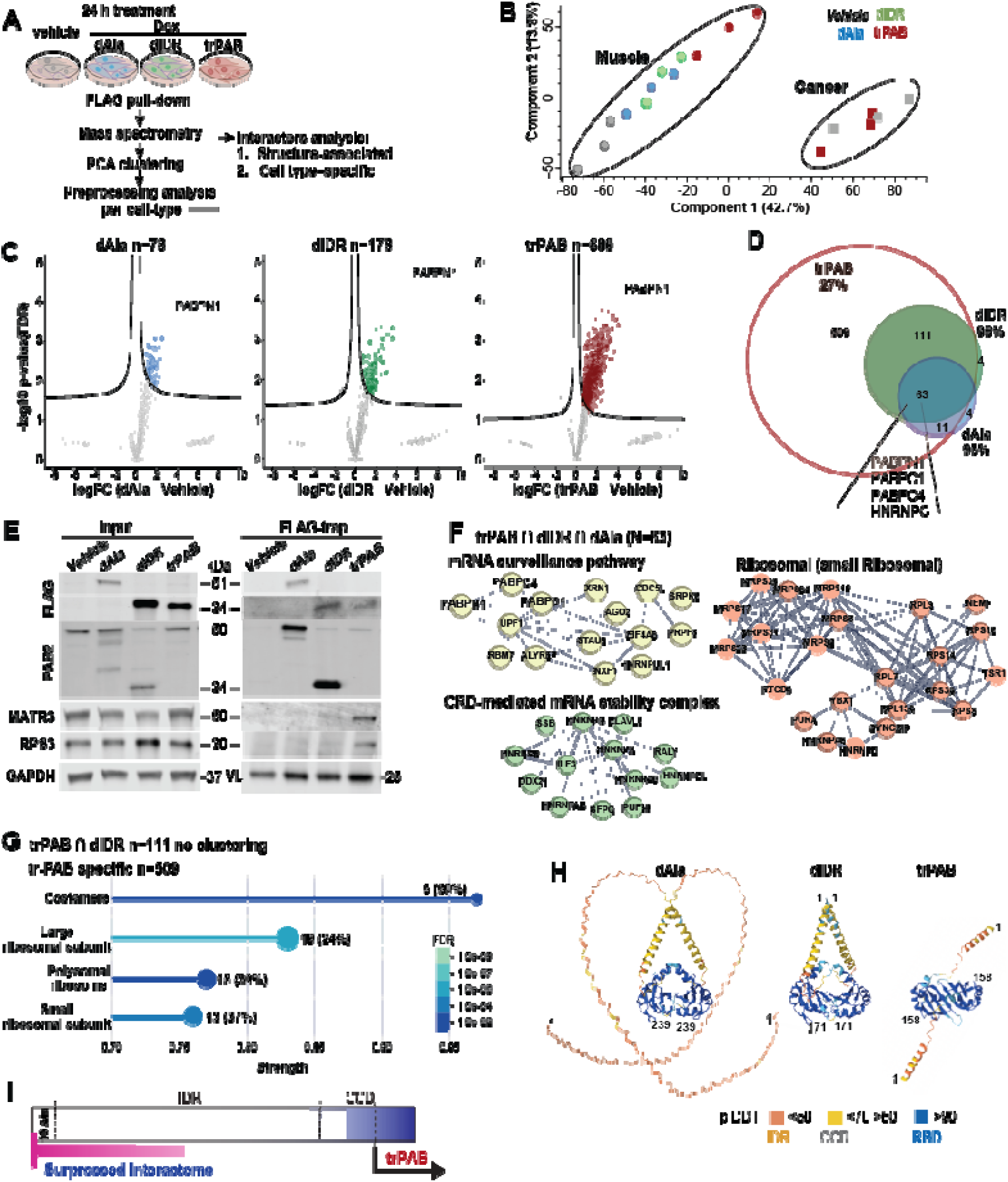
Interactome of PABPN1 N-terminal structural variants. **A.** Schematic overview of the experimental workflow for PABPN1 interactome analysis. **B**. PCA of mass spectrometry data from muscle and bladder cancer cells together. The major sources of variance (in %) are captured by PC1 and PC2. Panels C–G show analyses performed in muscle cells. **C**. Volcano plots showing differential interactors for each PABPN1 variant relative to vehicle control (log₂ fold change vs. –log₁₀ P value). Significant interactors (FDR < 0.05) are coloured and indicated (n=). Proteins selected for validation are labelled. The ‘wings’ under the S curves represent proteins that have values in 2 out of the three samples. **D.** Venn diagram showing overlap of interactors between PABPN1 variants. Literature-based PABPN1 interactors are depicted. **E.** Co-immunoprecipitation validation of selected proteins using FLAG-trap. Input controls show protein expression levels across conditions. GAPDH serves as a loading control for inputs, and variable light chain (VL) confirms immunoprecipitation. **F.** Protein-protein interaction clustering of the 63 shared interactors between dAla, dIDR, and trPAB. **G**. Clustering analysis results of the shared interactors between dIDR and trPAB (n = 111, no significant clusters), and a lollipop plot of c trPAB specific interactors (N = 509). The number of proteins and the enrichment % are depicted; colour indicates FDR-adjusted P value. **H.** AlphaFold3 structural predictions of dAla, dIDR, and trPAB variants modelled from N-terminus to the RBD. In all models the orientation of the RBD is horizontal. The positions of the first and last residues indicated. Predicted local distance difference test (pLDDT) scores are shown as a colour scale: blue (high confidence, RNA-binding domain), cyan to yellow (moderate to low confidence, coiledcoil domain), and orange (low confidence, intrinsically disordered region). **I.** A schematic summary of quantitative interactome and length-dependent regulation of N-terminal domains in PABPN1

Interactor abundance increased markedly with decreasing IDR length: 78 proteins were identified for dAla, approximately twofold more for dIDR (n = 178), and up to sevenfold more for trPAB (n = 689; Fig. 2C). Comparative analysis revealed extensive overlap between the trPAB interactome and those of dAla and dIDR (95% and 98%, respectively; Fig. 2D). A core set of 63 proteins was shared across all three variants, including PABPN1 and known interactors such as PABPC1, PABPC4, and HNRNPC, supporting the robustness of the dataset. Mass spectrometry findings were validated by immunoprecipitation and immunoblotting for FLAG, PABPN1, MATR3, RPS3, and IGF2BP2, showing the strongest co-immunoprecipitation signals in trPAB (Fig. 2E and Fig. S3A). Co-immunoprecipitation (IP) in A16 cells after 24 hours of doxycycline (Dox) induction showed efficient FLAG pull-down but no co-IP of PABPN1, MATR3, or RPS3 (Fig. S3B). Notably, these proteins were detected in the insoluble fraction of A16 cells ^18^, consistent with aggregation-associated loss of soluble interactions. Protein–protein interaction network analysis of the shared proteins (n = 63) revealed three major clusters: the mRNA surveillance pathway, CRD-mediated mRNA stability complexes, and ribosomal proteins enriched for the small subunit (Fig. 2F).

To identify strong PABPN1 binders, we applied hierarchical and k-means clustering to the fold-change values in muscle (n=9) and cancer (n=3) (Fig. S4A, B). A small protein cluster displayed expression patterns closely matching those of PABPN1 (Fig. S4B; n = 44 proteins), suggesting the presence of strong interactors. Among these were PABPC1 and PABPC4 (Fig. S4C). Protein–protein interaction network enrichment analysis of this cluster revealed enrichment for pathways related to nuclear mRNA processing, apoptosis-induced DNA fragmentation, and cytosolic translation, including the small ribosomal subunit and the coding region instability determinant (CRD)-mediated mRNA stability complex (Fig. S4C). These findings point to previously unexplored roles of PABPN1.

While the 111 proteins shared between trPAB and dIDR did not show significant clustering, the 501 trPAB-specific interactors were strongly enriched for ribosomal (small and large subunit) and polysomal proteins (Fig. 2G). Notably, myofiber structural proteins, costamere-associated components, were uniquely enriched in the trPAB interactome (Fig. 2G). These data indicate that the IDR constrains PABPN1 interactions in a length-dependent manner. Consistent with this, AlphaFold3 modeling of the N-terminus and RBD predicts that the IDR and CCD fold onto the RBD, whereas trPAB adopts a more open conformation (Fig. 2H). This supports a model in which the N-terminal region restricts access to the RBD, thereby limiting protein interactions (Fig. 2I). The trPAB interactome suggests increased accessibility to ribosomal proteins and enhanced engagement with the translation machinery and myofiber structural components. These findings raise the possibility that trPAB expression may restore PABPN1 function without promoting aggregation or cellular toxicity.

### The trPAB interactome in cancer cells is more limited than in muscle cells

trPAB colocalized with endogenous PABPN1 in the nucleus (Fig. 3A). In bladder cancer cells, trPAB also co-immunoprecipitated with PABPN1 (Fig. 3B). Principal component analysis showed clear separation of trPAB samples from vehicle-treated controls along PC1 (Fig. 3C), and 96 interactors were identified (Fig. 3D). PABPN1 was the most significantly enriched protein, with PABPC1 and PABPC4 also among the top interactors (Fig. 3D). Across cell types, 52 interactors were shared between bladder cancer and muscle cells, enriched for ribosomal proteins and negative regulators of mRNA metabolism (Fig. 3E). A cluster specific for PABPN1-interactome in cancer cells (n=44, and Fig. S4B) was enriched for mitochondrial translation (Fig. 3E). Together, these findings indicate that trPAB maintains a conserved core interactome while exhibiting cell type–specific interactions.

**Figure 3.**
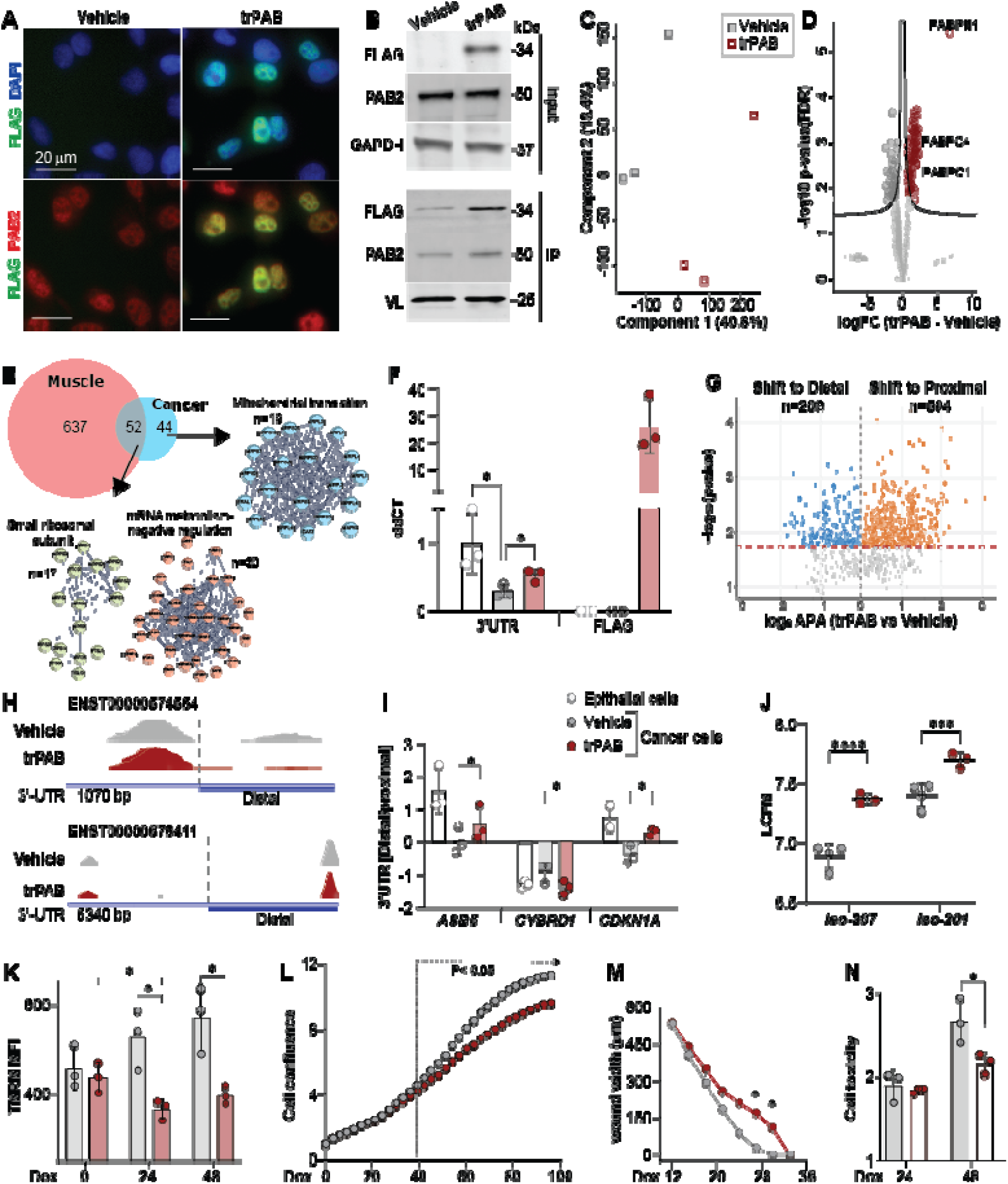
trPAB restores cellular function in bladder cancer cells. **A.** Immunofluorescence showing nuclear colocalization of trPAB (green) and PABPN1 (red) in bladder cancer cells. Nuclei are counterstained with DAPI (blue). Scale bar, 20 µm. **B.** Western blot of trPAB expression and FLAG immunoprecipitation (IP) following 24 h Dox induction. PABPN1 is detected using a PAB2 antibody. GAPDH serves as a loading control for input, and the variable light chain (VL) in the IP. Molecular weights are in kDa. **C-D.** PCA plot (C) and Volcano plot (D) of trPAB interactors vs. vehicle control (logC fold change vs. –logCC *P* value). Significant interactors (FDR < 0.05, FC ≥ 2) are highlighted in red. PABPN1, PABPC1 PABPC4 are highlighted. **E.** Venn diagram shows the overlap of trPAB interactors in muscle (orange) and bladder cancer (cyan) cells and the protein-protein interaction clustering of the 61 shared interactors (yellow and red) and the 81 cancer unique interactors (blue). **F.** RT–qPCR analysis of *PABPN1* expression (ΔΔCt) measured with primers targeting the 3′ UTR (endogenous) and FLAG (trPAB) in epithelial and bladder cancer cells treated with vehicle (v) or Dox for 24 h. **G.** Vulcano plot of APA-shift (p<0.05, FDR) in trPAB vs control. A shift to distal is in orange and to proximal in blue. The dotted red line is p=0.05 FDR. **H.** IGV plots of average reads at the 3’-UTR in vehicle (grey) and trPAB (red). The Distal half and the 3’-UTR length are denoted. **I.** RT–qPCR analysis of alternative polyadenylation measured using primer sets to distal and proximal sides of the 3’-UTR. A change in proximal/distal ratio shows APA-shift. **J.** Dot plot of PABPN1 reads count (log CPM) for isoforms 207 (Δexon-1) and 201 (full length). Read counts was from RNAseq. **K–O.** Functional assays in Dox- or vehicle-treated (grey) and trPAB–expressing (red) cells. Treatment time is in hours. **K.** Mitochondrial membrane potential (TMRM MFI). **L.** Cell growth (confluence over 96 h; significance from 39 h). **M.** Cell migration (wound closure over 32 h; significance at 27–30 h). **N.** Cell toxicity. Every dot represents a biological replicate; one way ANOVA was applied for two samples and two-way ANOVA for three samples both statistical analysis with a post-hoc Tukey test. *P* < 0.05 (*)*, < 0.01 (**), < 0.005 (***),* < 0.0001 (****).

### trPAB restores PABPN1 molecular function and cancer cell phenotypes

We next assessed whether trPAB restores PABPN1 levels in bladder cancer cells. Consistent with a previous study ^13^, we confirmed reduced *PABPN1* expression levels in bladder cancer cells compared with epithelial cells, while trPAB restored *PABPN1* levels (Fig. 3F). Reduced PABPN1 levels have been linked to altered polyadenylation site usage and alternative polyadenylation (APA) ^10,24^. The 1C library preparation robustly captures APA events at 3′-UTRs in muscle tissue and muscle cells ^10^. In cancer cells, trPAB leads to higher reads count at the 3’-UTR at both proximal and distal regions of 3′ UTRs (Fig. S5), from these 713 transcripts showing significant APA shifts (Fig. 3G). Changes in reads between proximal and distal sides of the 3’-UTR were then confirmed using IGV plots (representative examples in Fig. 3H). Additionally, RT–qPCR using primer sets targeting proximal and distal 3′ UTR regions showed that in trPAB expressing cells APA ratio of key cell cycle regulators is restored (Fig. 3I). Reads counts from RNAseq showed that the abundance of both PABPN1 isoform 207 (lacking exon 1 as in trPAB) and isoform 201 (the full-length *PABPN1*) was increased in trPAB-expressing cells (Fig. 3J). Together, these results show that trPAB restores PABPN1 levels and supports functional recovery of PABPN1 activity.

We then assessed energy metabolism using TMRM that determines mitochondrial membrane potential ^25^. Energy metabolism was higher in bladder cancer cells compared with epithelial cells (Fig. S6), which was returned by trPAB as early as 24 hours post-induction (Fig. 3K), suggesting that energy metabolism is restored by trPAB expression. In addition, trPAB expression following 72 hours Dox treatment reduced both cell proliferation and migration (Figs. 3L and 3M). Notably, the decrease in cell growth became evident approximately 40 hours after induction and coincided with reduced cellular toxicity (Fig. 3N). This temporal sequence suggests that the trPAB interactome influences APA, leading to subsequent improvements in mitochondrial function, which ultimately translate into changes in cellular phenotypes such as proliferation and migration.

### PABPN1 variant trPAB improves muscle cell features

We next assessed the effect of trPAB on muscle cell differentiation using immunofluorescence for myosin heavy chain (MyHC), a marker of fusion and myofiber maturation ^26^. Increased MyHC signal in trPAB-treated cultures indicates greater cell fusion compared with vehicle-treated controls (Fig. 4A-B). Moreover, the area of multinucleated cells was larger in trPAB-treated cultures, suggesting enhanced formation of multinucleated myotubes (Fig. 4C). Consistent with this, MYOG signal, a myogenic transcription factor regulator of cell fusion ^27^, was elevated in differentiated cultures and within multinucleated cells (Fig. 4D-E), indicating activation of the myogenic program.

**Figure 4.**
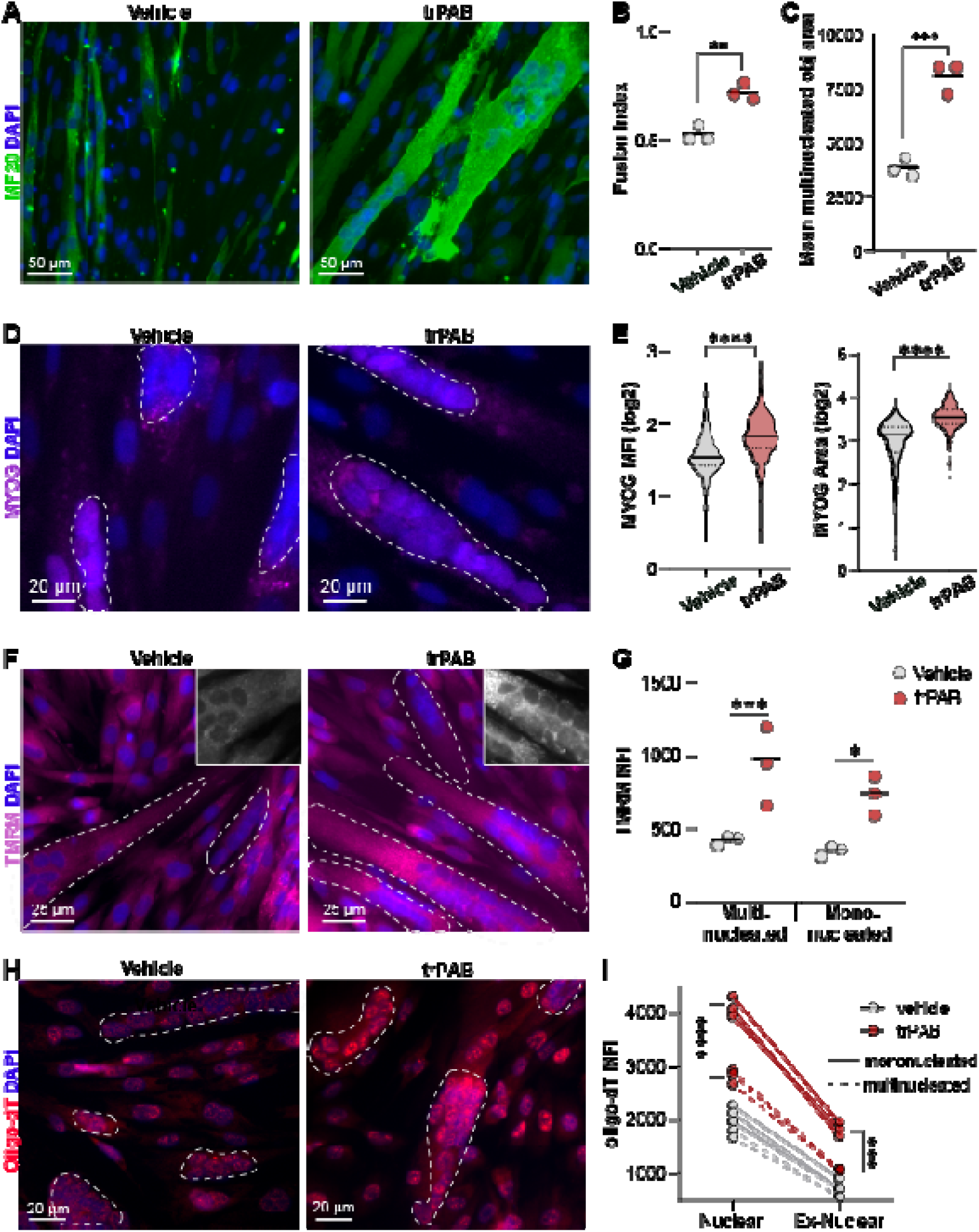
trPAB improves muscle cell function. **A.** Representative immunofluorescence images of differentiating vehicle-treated or trPAB cultures stained with MF20 (green); nuclei in blue. Multinucleated cells are encircled with a dashed line. Scale bar, 50 µm. **B-C.** Analysis fusion index (B) or multinucleated object area (C). Each dot represents one biological replicate (mean of 3,000–4,000 cells). One-way ANOVA. **D.** Representative immunofluorescence images of MYOG staining (red) and nuclei in blue. The multinucleated cells are encircled with a dashed line. Scale bar, 20 µm. **D.** Analysis of MYOG fluorescence intensity (left) or MYOG object area (right). Each violin plot represents mean of 600-1000 cells). One-way ANOVA. **F.** Representative images of TMRM staining in differentiating cultures; nuclei in blue, the insert shows TMRM channel. Multinucleated cells are encircled with a dashed line. Scale bar, 25 µm. **G.** Dot plot of mean TMRM signal per object in multinucleated and mononucleated objects, each dot represents one biological replicate (mean of 3,000–4,000 cells). Two-way ANOVA. **H.** Representative images of oligo(dT) in situ hybridization (red), the nuclei in blue. Multinucleated cells are encircled with a dashed line. Scale bar, 20 µm. **I.** Analysis of mean oligo(dT) signal per mono-nucleated or multinucleated objects in nuclear or perinuclear compartments. Each dot represents one biological replicate (mean of 3,000–4,000 cells). One-way ANOVA with a post-hoc Tukey test. *P* < 0.05 (*)*, < 0.01 (**), < 0.005 (***),* < 0.0001 (****).

Given the role of PABPN1 in alternative polyadenylation (APA) and mRNA processing, we examined downstream functional outputs. In trPAB cells the mitochondrial activity was increased, with a more pronounced effect in multinucleated cells (Fig. 4F, G). Moreover, oligo(dT) in situ hybridization in differentiated cells revealed increased poly(A)+ RNA signal in both nuclear and cytoplasmic compartments (Fig. 4H and 4I), suggesting improved mRNA metabolism: nuclear export, and stability. Together, these findings suggest that trPAB enhances cytoplasmic mRNA availability that ultimately promote efficient myogenic differentiation. Notably, in bladder cancer cells trPAB did not affect mRNA accumulation and subcellular accumulation (Fig. S7), further strengthen cell type specific activity of trPAB.

### trPAB restores muscle cell defects driven by the expression of the expanded PABPN1

Overexpression of full-length PABPN1 in muscle cells induces aggregation, and the expanded PABPN1 variant (A16) impair muscle cell function, including reduced fusion capacity and mitochondrial activity, in association with genome-wide proximal APA ^18^. We therefore assessed whether trPAB could rescue A16-driven defects in differentiated muscle cells using a co-culture approach (Fig. 5A). Stable A16 and trPAB muscle cells were co-cultured and induced to differentiate. Dox was applied either prior to differentiation to assess fusion, or in differentiated cultures to evaluate trPAB effects (Fig. 5A). In co-cultures of A16-YFP and trPAB, multi-nucleated myotubes contained nuclei positive for either YFP or FLAG (Fig. 5B). Notably, the fraction of FLAG-positive nuclei was reduced in co-culture compared to trPAB-only cultures (Fig. 5C), while A16 puncta area and intensity were significantly decreased upon trPAB co-expression (Fig. 5D), indicating modulation of A16 aggregation.

**Figure 5.**
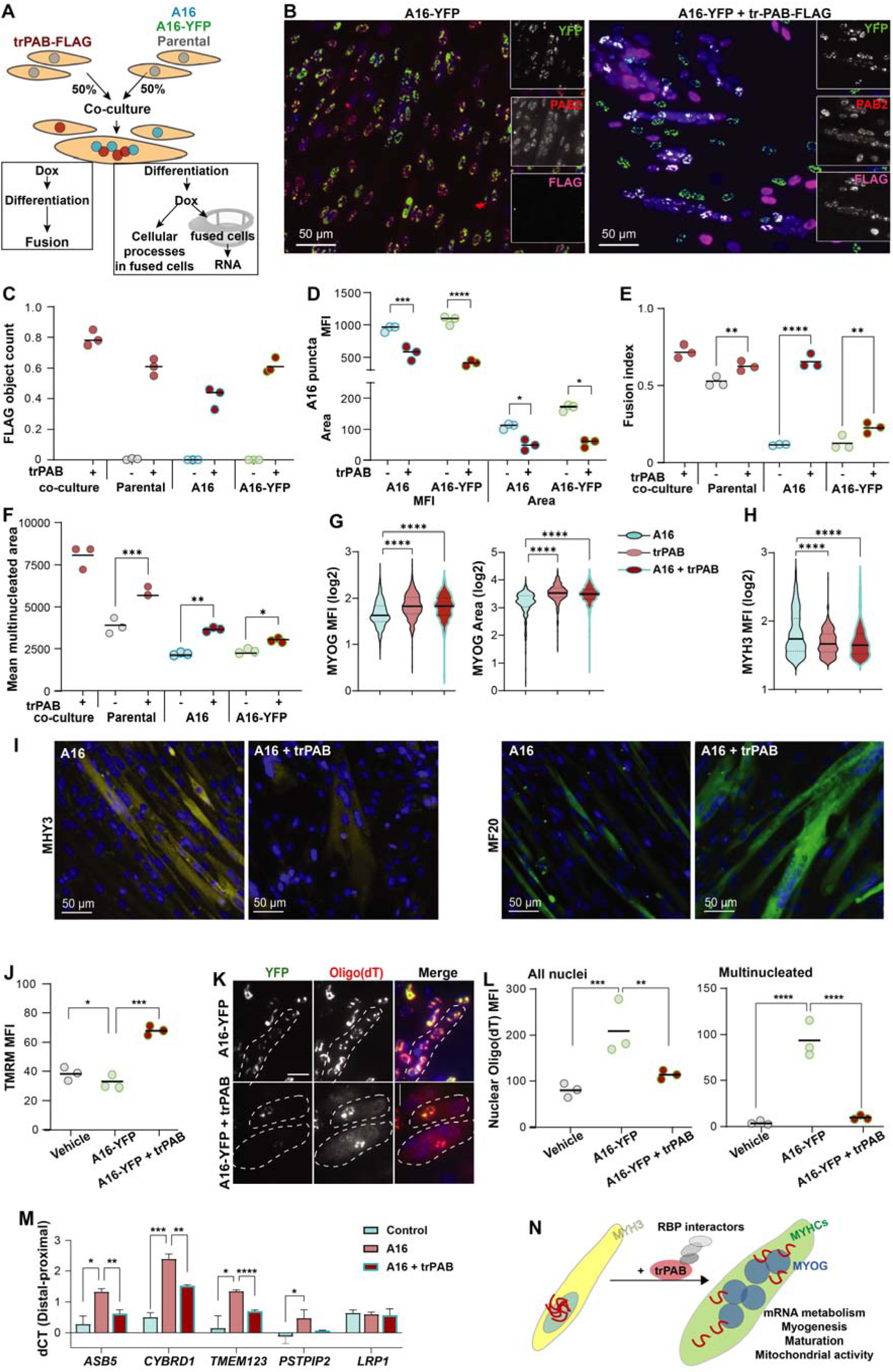
The PABPN1 variant, trPAB, restores A16-driven differentiation and RNA processing in co-culture systems. **A.** A schematic overview of the co-culture experimental design. **B.** Representative fluorescence images of A16-YFP cells (left) and A16-YFP co-cultured with trPAB-FLAG. Insets show single-channel images (YFP, PABPN1, FLAG) from fused multinucleated cells. Scale bar, 50 µm. **C–F.** Quantification of co-culture experiments with trPAB-FLAG in A16-YFP, A16, or parental cells. trPAB alone serves as reference. Cells were Dox-treated during differentiation. Each dot represents one biological replicate (mean of 3,000–4,000 cells). Two-way ANOVA. (**C)** Fraction of FLAG-positive nuclei. (**D)** Puncta area and mean fluorescence intensity (MFI) of A16-YFP or PABPN1 (PAB2). (**E)** Fusion index (fraction of nuclei within MF20-positive structures). (**F)** Mean multinucleated area. Two-way ANOVA with a post-hoc Tukey test (D-F). **G–H.** Violin plots of differentiation markers in multinucleated cells. (**G)** MYOG MFI and area. (**H)** MYH3 MFI. Data represent 700–1,000 multinucleated objects. One-way ANOVA with a post-hoc Tukey test. (G-H). **I.** Representative immunofluorescence images of differentiating A16 or A16 + trPAB-FLAG cultures stained for MYH3 (yellow) and MF20 (green); nuclei in blue. Scale bar, 50 µm. **J–N.** Analysis of differentiated A16 and A16 + trPAB-FLAG cultures following 24 h Dox treatment; vehicle-treated cells serve as reference. Each dot represents a biological replicate (1,000–1,500 cells). (**J)** Mitochondrial membrane potential (TMRM MFI); One-way ANOVA with a post-hoc Tukey test. **K–L.** Oligo(dT) in situ hybridization. (**K)** Representative images showing oligo(dT) and YFP staining; multinucleated cells outlined (dashed line). Scale bar, 20 µm. (**L)** Quantification of oligo(dT) MFI in mononucleated (left) and multinucleated (right) cells. Two-way ANOVA with a post-hoc Tukey test. **M–N.** RNA analysis in differentiated cells (mean ± SD, n = 3). One-way ANOVA with a post-hoc Tukey test. **M.** Alternative polyadenylation (APA) measured as distal-to-proximal usage ratio (ΔCt) for *ASB5*, *CYBRD1*, *TMEM123*, and *PSTPIP2*; *LRP1* as control. *P* < 0.05 (*)*, < 0.01 (**), < 0.005 (***),* < 0.0001 (****). **N.** Schematic model summarizing layers of trPAB effect in muscle cell differentiation.

Functionally, trPAB significantly improved the fusion index and increased the size of multinucleated myotubes in A16 cultures (Fig. 5E and 5F). Myogenesis is marked by transient expression of myogenic transcription factors, including MYOG ^28^, and the expression of MYH3 mark immature fibers ^29^, whereas mature MYHC isoforms are labelled with MF20 ^30^. MYOG signal was elevated in myofibres from the trPAB and A16–trPAB co-cultures compared with A16 cultures, whereas MYH3 signal was higher in the A16 culture (Fig. 5G–H). MYH3-positive myofibres were more prevalent in A16 cultures than in A16–trPAB co-cultures, while MF20-positive myofibres were more abundant in A16–trPAB co-cultures (Fig. 5I), indicating enhanced progression toward mature myofibres. Mitochondrial activity was also enhanced in A16–trPAB myotubes (Fig. 5J).

At the molecular level, A16 myotubes exhibit nuclear retention of poly(A)+ RNA co-localizing with PABPN1 aggregates ^18^. In A16–trPAB co-cultures, nuclear mRNA accumulation was reduced, particularly in multinucleated myotubes (Fig. 5K and 5L), suggesting restoration of mRNA export and metabolism. We selected four genes from the A16 APA-shift study ^18^ to assess if APA-shift is restored in A16-trPAB myotubes. RT-qPCR analysis showed reversal of A16-associated proximal APA shifts, whereas unaffected transcripts (*LRP1*) remained unchanged (Fig. 5M). Together, these data support a model in which trPAB counteracts A16-induced aggregation and restores RBP interactions, leading to normalization of APA and mRNA metabolism (Fig. 5N). This, in turn, enhances, mitochondrial function and myotube maturation, thereby rescuing key aspects of muscle cell pathology.

### trPAB reverses histopathological phenotypes in an OPMD mouse model

Our findings suggest that trPAB may represent a therapeutic strategy for OPMD. To test this in vivo, we delivered trPAB-FLAG via an AAV9 vector into tibialis anterior (TA) muscles of the A17.1 OPMD mouse model (hereafter A17) (Fig. 6A). Saline-injected A17 and wild-type FVB mice served as controls. Western blot analysis confirmed trPAB expression in all injected muscles, albeit with variable levels across animals, and showed increased total PABPN1 levels (Fig. 6B and 6C).

**Figure 6.**
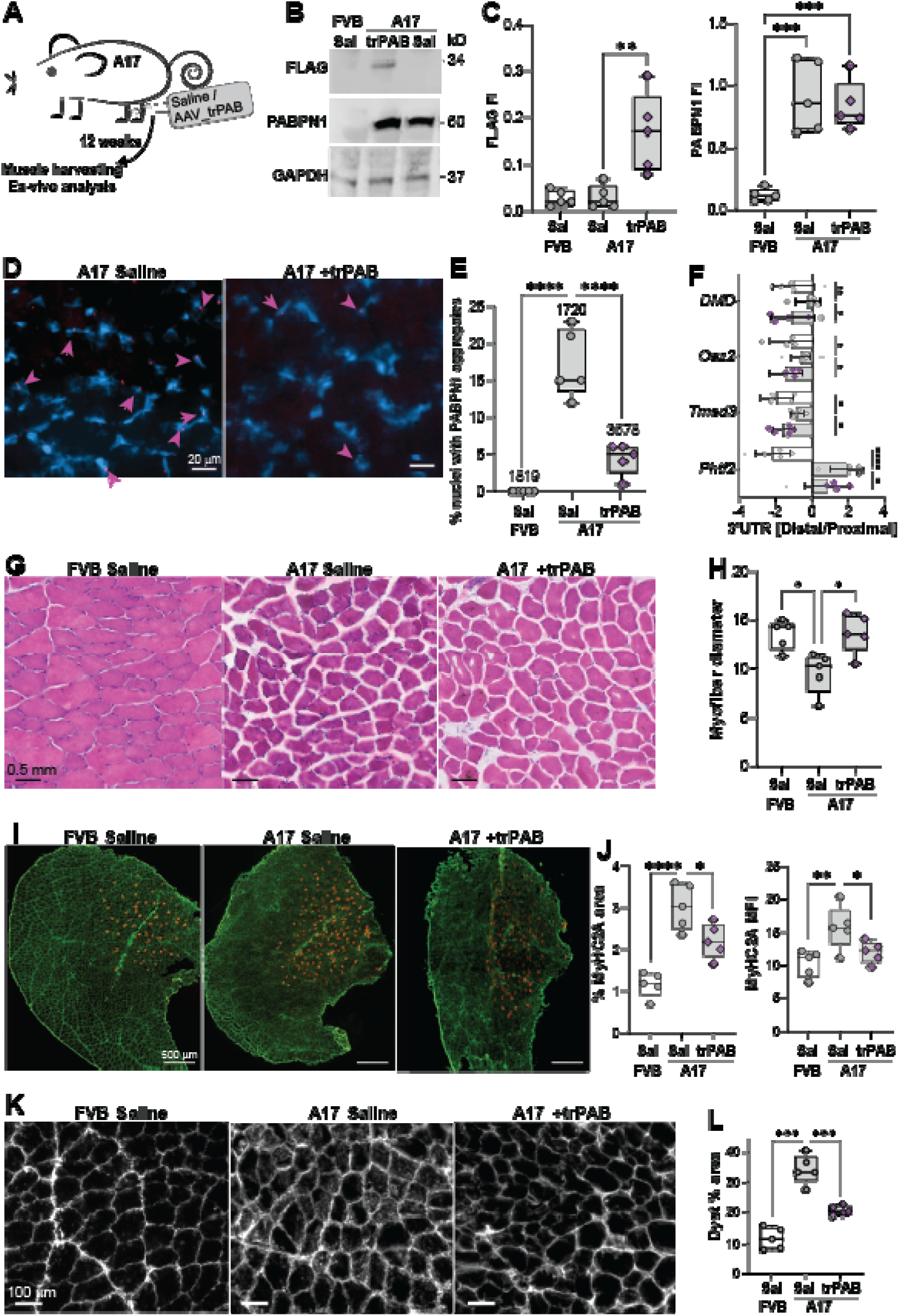
trPAB restores muscle histopathology in A17.1 TA muscle. Analysis was performed in saline-treated FVB and A17.1 (A17) mice, and in AAV9–trPAB–treated muscles (n = 5 mice per group). **A.** Schematic overview of the in vivo treatment. **B.** Representative Western blot showing FLAG, PABPN1 (PAB2), and GAPDH. **C.** Quantification of FLAG and PABPN1 fluorescence intensity (FI). **D.** Representative images of aggregated PABPN1 (red); arrows indicate nuclei with aggregates in A17 saline- and trPAB–treated muscles. Scale bar, 20 µm. **E.** Percentage of nuclei containing aggregated PABPN1. Total nuclei analyzed are indicated. **F.** Alternative polyadenylation (APA) measured as distal-to-proximal usage ratio (ΔCt) for *DMD*, *Oaz2*, *Tmed3*, and *Phtf2*. **G.** Representative HE images. Scale bar, 0.5 mm. **H.** Quantification of mean myofiber diameter. Each dot represents the average of 400-600 myofibers from one mouse. **I.** Representative immunofluorescence images of MyHC-2A (red) and Dystrophin (green) across entire muscle cross-sections. Scale bar, 500 µm. **J.** Quantification of MyHC-2A area (left) and mean fluorescence intensity (MFI, right). **K.** Representative immunofluorescence images of Dystrophin. Scale bar, 100 µm. **L.** Quantification of Dystrophin-positive area. All data are presented as box plots. One-way ANOVA Dunn’s correction for multiple comparisons: *P* < 0.05 (*)*, < 0.01 (**), < 0.005 (***),* < 0.0001 (****).

Nuclear PABPN1 aggregation is a hallmark of A17 pathology. Using a PAB2 antibody that does not recognize trPAB, we detected A17-associated PABPN1 aggregates and observed a significant reduction in the proportion of aggregate-containing nuclei in trPAB-treated muscles compared to saline-injected A17 controls (Fig. 6D and 6E), consistent with our cell-based findings.

At the molecular level, trPAB restored alternative polyadenylation (APA) at the 3′-UTR of all four genes tested, DMD, Oaz2, Tmed3, and Ohtf2 (Fig. 6F), whose APA is disrupted in A17 mice ^10,31^. We previously reported APA events within *DMD* intragenic regions ^32^, indicating the presence of alternatively spliced protein variants that may give rise to distinct Dystrophin isoforms. Such variants have been reported to exhibit intracellular accumulation, rather than the predominantly sarcolemmal localization ^33^.

A17 myofibers are atrophic and display elevated expression of MyHC-2A ^34^. Treatment with trPAB restored myofiber diameter (Fig. 6G–H) and MyHC-2A fluorescence intensity and area (Fig. 6I–J), indicating reversion of atrophy. These results suggest that while trPAB effectively rescues myofiber atrophy, a measurable increase in overall muscle mass may require longer treatment duration.

In addition, A17 muscles showed abnormal cytosolic localization of dystrophin alongside its typical membrane distribution. Quantification revealed an increased fluorescence area compared to FVB controls (Fig. 6G–H). Cytosolic dystrophin has been reported in immature muscle fibers ^35^, consistent with MYH3 expression observed in the A16 cell model. trPAB treatment partially restored proper dystrophin localization and reduced the aberrant fluorescence area (Fig. 6G–H), suggesting improved dystrophin organization, myofibre maturation and function.

Collectively, these findings demonstrate that trPAB reduces pathogenic PABPN1 aggregation, restores APA-dependent RNA processing, and ameliorates key histopathological features of OPMD muscle. This supports improved muscle fibre integrity and highlights the therapeutic potential of this approach.

## Discussion

In this study, we explored the effect of structural regions at the N-terminus of PABPN1 on its protein features, aggregation and function. Surprisingly, the short coiled-coiled domain contains two functionally distinct regions: the amino acids at the N-terminus are critical for PABPN1 stability, whereas the second half of the CCD plays a different role by limiting aggregation. The CCD is generally associated with structural rigidity and protein interactions ^36^. In PABPN1 the CCD has been implicated as a structural element promoting PABPN1 aggregation ^19–21^. These studies consider the CDD as a single functional domain associated with its structural rigidity. Our data refine this view by demonstrating that protein stability and aggregation are regulated by different amino acid sequences, indicating two discrete functional domains of the alpha-helix structure.

We further found that the intrinsically disordered region (IDR), located between the first methionine and the CCD, plays a key role in shaping the PABPN1 interactome. Consistent with its disordered nature and AlphaFold predictions, this IDR lacks a fixed structure. Despite this, our interactome analysis indicates that the IDR acts as a gatekeeper: its presence, and particularly its length, reduces the number of interacting partners. This effect is even more pronounced in the pathogenic form of PABPN1 carrying an expanded alanine tract, which also shows reduced self-interaction. IDRs are widespread across diverse proteins and are thought to play central roles in essential cellular processes ^37–39^. Their functions have primarily been studied through experimental and computational analyses of their conformational ensemble ^37^. These ensemble properties can influence protein folding, binding to other proteins, and the spatial organization of protein complexes ^37^. The IDR in PABPN1 regulates aggregation and shapes the PABPN1 interactome in a cell type-dependent manner. In muscle cells, IDRs are often linked to phase separation and aggregation ^38^. Our results show that removal of the IDR reduce aggregation, while removal of part of the CCD prevents aggregation, indicating that the IDR contributes to, but does not solely drive, aggregation. Moreover, aggregation is strongly influenced by cellular context. In muscle cells, aggregation occurs rapidly, within 24 hours, whereas in bladder cancer cells, it is not observed under the same conditions. Moreover, in muscle cells aggregation differs between proliferating and differentiating cells ^22^. This highlights that PABPN1 aggregation depends not only on intrinsic protein features but also on the cellular environment. Although IDRs are generally associated with flexible binding and interaction diversity ^37^, the IDR of PABPN1 appears to constrain the interactome in a cell type-dependent manner. Particularly interactions with RNA-binding proteins are supressed, suggesting modulation of PABPN1 activity. Comparison of the trPAB and IDR-deleted interactomes suggests that the first half of the CCD also restricts protein interactions, possibly by imposing a more rigid conformation that limits accessibility. Analysis of binding patterns across the four conditions suggests that in addition to its known nuclear function, PABPN1 could be involved in translation and mediation of mRNA stability during translation, agreeing with publications reporting PABPN1 effect on these cytosolic functions ^15,40^ In cancer cells, PABPN1 might be involved in mitochondrial translation, suggesting that PABPN1 acts in a cell-type specific manner.

We identify trPAB, a naturally occurring PABPN1 variant, as a stable, non-aggregating form of the protein. Functionally, trPAB enhances myogenesis. In muscle cells, but not in bladder cancer cells, the trPAB interactome is enriched for costamere-associated proteins. In addition, proteins involved in the translation machinery are strongly enriched, consistent with previous work showing that PABPN1 regulates cytoplasmic mRNA processing and translation ^40^. In line with this, reduced PABPN1 levels impair translation, particularly at the level of polysomes. Here, we further show that trPAB expression increases cytoplasmic mRNA accumulation in muscle cells. This effect is context-dependent, as it is not observed in bladder cancer cells and is more pronounced in differentiated muscle cells. Together, these findings indicate that trPAB function is strongly shaped by cellular context and is particularly relevant to muscle biology, supporting a protective or cellular profile.

In cell models with reduced PABPN1 function, including bladder cancer cells and A16 muscle cells, trPAB overexpression reverses the pathogenic phenotype without detectable toxicity. In vivo, trPAB delivered via AAV9 to tibialis anterior muscle in an OPMD model reduces aggregation, restores PABPN1 function, and improves myofiber pathology. A two-step “silence-and-replace” strategy is emerging as a therapeutic approach for Oculopharyngeal Muscular Dystrophy, in which endogenous PABPN1 (both wild-type and mutant) is silenced and replaced with a codon-optimized variant ^41^, https://musculardystrophynews.com/bb-301/. While effective, this two-step approach is complex. Our findings suggest that using a naturally occurring variant such as trPAB may provide a simpler and potentially safer alternative.

Together, our results identify N-terminal regulation of PABPN1 as a key determinant of RNA processing capacity. In muscle cells, but not in bladder cancer cells, we link protein aggregation to alternative polyadenylation (APA)-dependent control of cellular function, revealing a tractable mechanism for therapeutic intervention. In contrast, in cancer cells, APA-related effects appear to be independent of PABPN1 aggregation.

## Methods

### PABPN1 constructs and virus production

*Generation of lentivirus constructs* - The deletion forms of PABPN1, and trPAB were made in the pCW57-MCS1-2A-MCS2 doxycycline (Dox) inducible lentiviral vector (#71782, Addgene) from the A16 expression vector ^18^ using the QuikChange Site-Directed Mutagenesis Kit (#200519, Agilent technology). Primer sets are in Table S1. All deletion and trPAB forms were confirmed by Sanger sequencing. Lentivirus production was performed as detailed in ^42^.

*Generation of AAV constructs* - thPAB-FLAG gene sequence with Kozak sequence was ordered from Genscript in pUC57 backbone and subcloned into a pAAV-HSA vector downstream of HSA promoter using the restriction enzymes EcoRI and MluI, generating pAAV-HSA-trPAB. AAV9 particles were prepared using a standard double transfection protocol as detailed in ^11^. In brief, HEK293T/C17 cells were seeded in roller bottles and cultured in Dulbecco’s modified Eagle’s medium (DMEM) containing 10% fetal bovine serum at 37°C and 5% CO2. Once 50% confluent, media was changed to DMEM containing 2% fetal bovine serum. Cells were then transfected with the relative pAAV plasmid and pDP9rs AAV serotype 9 helper plasmids using polyethylenimine at a ratio of 1:4. Following 72 hours incubation, cells were lysed and viral particles were precipitated from supernatant using polyethylene glycol-2000. Viral particles were then purified using iodixanol (Sigma-Aldrich, Gillingham, UK) step-gradient ultracentrifugation, desalted and concentrated. The copy number of vector genomes was quantified by quantitative polymerase chain reaction with primers to trPAB (Table S1).

### Mouse

A17.1 male mice and FVB, WT, age and sex matched were used. Animals were housed with food and water *ad libitum* in minimal disease facilities (Royal Holloway, University of London)*. In vivo* experiments were conducted under statutory Home Office recommendation; regulatory, ethics, and licensing procedures and the Animals (Scientific Procedures) Act 1986 (Project Licence PP60551761).

*AAV delivery* - Twelve-week-old A17.1 or FvB mice were anesthetised with isoflurane, and 5x10^10^ vp of AAV-HSA-trPAB, diluted in 30 μl saline, were intramuscularly administered into *tibialis anterior* (TA) muscles (n=5 per group). Saline injected TA of both A17.1 mice and FvB mice, were used as negative and healthy controls respectively. Twelve-weeks post-injection, *in situ* TA muscle physiology was performed under terminal anaesthesia. Mice were weighted prior to injections and prior to sacrifice. After sacrifice, TA muscles were harvested, weighed, and either blocked in OCT and frozen in liquid nitrogen-cooled isopentane or snap froze in liquid nitrogen depending on the assessment to perform. For staining, the nitrogen-cooled isopentane were used.

### Cell culture

#### Muscle cell propagation

The immortalized human muscle clone (MB135) and growth conditions were as described in ^18^. The A16 and A16-YFP stable cells were described in ^18,22^, respectively. Myoblasts were cultured in growth medium (F10 (Gibco) medium supplemented with 15% FBS, 1 ng/ml bFGF-2 (#145aa, Canvax), and 0.4 µg/ml Dexamethasone (#D4902, Sigma-Aldrich). Cells were passaged when reaching 50-80%, and cell cultures were renewed every 6-8 weeks. Cell cultures did not reach 100% confluence to avoid spontaneous differentiation. Cells were frequently monitored for mycoplasm.

*Muscle cell differentiation* was done at high confluency (85-95%) in DMEM+2% horse serum for 3-5 days.

#### Cancer cell propagation

Bladder cancer cells, UMUC-3 (#CRL-1749, ATCC), and Immortalized Normal Human Urothelial Cells, NHU-Tert (#T0785, ABM) were cultured in DMEM supplemented with 10% FBS, 100 units/ml penicillin, 50 μg/ml streptomycin. Cells were passage 1:10. Cells were frequently monitored for mycoplasm.

#### Generation of stable cells

*Muscle* or cancer cells were incubated with lentivirus particles + polybrene as detailed in ^42^. Stable cells were selected with 2.5 mg/ml puromycin (#P7225, Sigma-Aldrich). The transgene was induced with 4 μg/mL doxycycline (Dox) (#D5207, Sigma Aldrich), and DMSO (1:1000) was used for (uninduced) vehicle-treated cells. Protein expression levels of all PABPN1 variants remained stable between 24 and 72 hours following Dox induction (Fig. S8). Therefore, unless otherwise specified, all experiments were conducted using a 24-hour Dox treatment.

#### Cell seeding for high content screening (HCS)

cells were seeded in a Nunc 96 well plate. For experiments in proliferating cells, cells were seeded at 40% confluence (20,000 cells per well), and differentiation cells were seeded at 90% confluence (50,000 cells per well). In muscle cells, Dox was added together with a differentiation medium or 3 days after differentiation medium for 24 hours.

#### Protein assays

*Protein extraction* – for all analysis was made with a RIPA lysis buffer (10 mM Tris pH 7.5, 150 mM NaCl, 0.5 mM EDTA, 0.1 % SDS, 1 % Triton™ X-100, 0.5% Tween- 20) and freshly added protease inhibitor cocktail (#A32955, ThermoFisher). After ice incubation for 30 minutes and vortex every 10 minutes, the supernatant was sonicated and collected after 10 min centrifugation, 17000g, at 4°C. Protein aliquots 30-40 mg were separated on SDS-PAGE gradient gel.

*Western blot* - was carried out with a PVDF membrane. Bulk proteins were visualized with the No-Stain Protein Labeling Reagent (#A44717, ThermoFisher) and imaged using the iBright Imaging System (ThermoFisher). The membrane was blocked with 5% dried milk powder (#T145.2, Carl Roth). Primary antibody incubation was carried out at 4°C overnight, and secondary antibody incubation at room temperature for one hour. Antibodies are listed in Table S2. The fluorescent signals on blots were scanned with an Odyssey CLx Infrared imaging system (LiCOR, NE. USA). Western blots from all mice are in Fig. S9.

#### Western blot quantification was done using ImageJ

Values were corrected for background and normalized for loading controls. Normalization was made for the No-Stain or housekeeping signal.

*FLAG-trap and immunoprecipitation* - Protein lysates for FLAG-trap experiments were prepared in RIPA buffer supplemented with 100 U DNase I (#EN0521, Thermo Fisher) and 2.5 mM MgClC, without sonication. To reduce nonspecific binding, Sepharose Protein A beads (Merck, #GE17-1279-01) were incubated with 1 mg protein lysate for 1 hour at 4°C (pre-clearing step). Following centrifugation at 17,000 × g for 10 min at 4°C, the unbound fraction was collected, and 700 µg protein was used for the FLAG-trap protocol (ChromoTek DYKDDDDK Fab-Trap® Agarose, #ffa, Proteintech) according to the manufacturer’s instructions. Briefly, beads were pre- washed and incubated with lysate for 1 hour at 4°C under rotation. Beads were then washed three times with RIPA buffer by centrifugation at 2,500 × g for 5 min at 4°C, followed by a final wash with PBS. Five percent of the beads were reserved for western blot analysis, and the remaining 95% were used for mass spectrometry. For western blot analysis, beads were incubated in 4× Laemmli Sample Buffer (#1610747, Bio-Rad) at 95°C for 5 min. For mass spectrometry, dry beads were subjected to an on-bead digestion protocol as described below.

#### Mass spectrometry and data analysis

Samples were subjected to an on-bead Smart digestion (Thermo Scientific) using 1 µL trypsin solution per sample, with digestion performed at 70°C, 1,400 rpm for 2 h. Tryptic peptides were desalted using the SOLA SPE (Thermo Scientific), dried by vacuum centrifugation and resuspended in 0.1% formic acid prior to MS analysis.

Tryptic peptides were analysed by LC-MS/MS using the VanquishNeo UHPL connected to an Orbitrap Ascend mass spectrometer with minor changes on the LC method from previously described method ^43^. Briefly, The Vanquish Neo was operated in “Trap and Elute” mode using an Acclaim PepMap trap (100um x 2cm) and EASY-SPRAY PepMapNeo column (50cmx75um, 1500bar). Tryptic peptides were trap and separated using a 60 min linear gradient (from 2 to 18 % B -100% ACN, 0.1% FA- in 40 min and from 18 to 35B% in 20 min at 300nl/min flow). MS data were acquired in Data-Independent acquisition mode (DIA), with 40 scan windows with variable width, as previously described ^43,44^.

Raw MS files were searched in three separate modes: one for all samples together (n = 18) and two searchers per cell-type: muscle (n=12) and bladder cancer cells (n=6) in DIA-NN v2.3.1 against a predicted library generated previously from the UniProt human proteome database (downloaded 5th Jan 2026), plus common contaminants, selecting Trypsin/P as enzyme and allowing one missed cleavage. Both MS1 and MS2 mass accuracy settings were inferred automatically from the data, and match-between-runs was enabled. Data pre-processing was carried out as follow: missing values replaces with zero, log2(x+1) transformation of reads count, exclusion of proteins with reads in only one sample per condition. Specific binders were identified against vehicle-treated cells, in a cell-type based analysis using with the FDR t-test (FDR < 5%) S0 0.1 in Perseus Software v2.1.6.0. The specific binders are listed in Table S3.

#### RNA extraction, library preparation, RNA sequencing, and analysis

RNA was extracted from vehicle (N=4) and 24 hours Dox-treated trPAB bladder cancer cells (N=3) using the PureLink RNA Mini Kit (Invitrogen 12183018A), according to the manufacturer protocol, continue with on-column DNase treatment PureLink™ DNase Set (Invitrogen 12185010). RNAs were stored at -80. RNA integrity was quantified using Qubit and checked on an RNA 6000 Nano Agilent Lab-on-a-Chip kit with Bioanalyzer Systems before cDNA library preparation. One hundred nanograms of RNA per sample were used to prepare poly(A) libraries with a single PCR cycle and 3’-UTR enrichment as detailed in ^45^. Sequencing was carried out with the Illumina NovaSeq 6000 system.

All reads in FASTQ format were first filtered using *Cutadapt* (v2.10), removing all remaining adapter sequences. *MultiQC* program in Python was used for quality control (QC) assessment of FASTQ files. The remaining reads were aligned to the Ensembl transcriptome version 104 using STAR (v2.7.5a), including UMI-based deduplication using UMI-Tools (v1.1.1), generating a transcriptome-based alignment in BAM format. We used the same Ensembl transcript annotation version 104 to create a customized transcript annotation GTF file for all Ensembl annotated transcripts. With the annotation and human transcriptome-based alignment files as input, we quantified the reads using *featureCounts* (v2.0.1). Only reads mapping to the 3′-UTR of transcripts were included in the analysis. Significant differentially expressed transcripts using the *t*-test (*FDR* <0.05; Table S4) were selected for the APA-shift calculation. APA-shift calculation is described in ^45^. With this approach, false positives arising from short 3′-UTRs were minimized and were insignificant. Transcripts with significant APA-shift (FDR <0.05) are listed in Table S4. Candidates were validated using IGV version 2.19.5. All analyses were performed using RStudio Software RStudio 2022.02.3 (Build 492) using R Statistical Software (v4.2.3).

#### RT-qPCR

RT-qPCR was conducted on RNA from the Dox-induced and vehicle-treated cells. RNA was extracted with the PureLink RNA Mini Kit (Invitrogen 12183018A). 500 ng RNA was reverse transcribed for cDNA synthesis using the QuantiTect Reverse Transcription Kit (QIAGEN) and random primers, following the manufacturer’s instructions. Subsequently, qPCR amplification was performed with the QuantiNova SYBR Green kit (QIAGEN) using 5 ng cDNA, with technical duplicates, using a standard amplification protocol at a melting temperature of 60°C. The PCR detection was carried out in the CFX384 system (BioRad). Samples with CT values above 35 were excluded from the analysis to eliminate potential noise. The average CT values from the technical duplicates and normalization to the HPRT1 gene were used for ddCT calculation. Primer sets were designed with the NCBI Primer design tool (https://www.ncbi.nlm.nih.gov/tools/primer-blast), and the primers are listed in Table S1.

#### Immunofluorescence and staining

*Cell culture –* cells were cultured in plastic bottom 96-well plates (#655101, Greiner™ 96-Well F-bottom Microplates). Cell fixation was performed with 4% Formaldehyde in PBS for 5 minutes. Subsequently, permeabilization was performed with 1% Triton-X100 for 10 minutes, followed by 1X PBS washing and first antibody incubation for one hour at room temperature. The first antibody was washed with excess (PBT PBS+0.05%-Triton-X) for five minutes each. Incubation with a fluorophore-conjugated secondary antibody and DAPI was carried out for 30 minutes, followed by PBT washing (3-times 5 minutes each). Cells were kept in PBS during imaging.

*Mouse* – staining was made in fresh cryosections. Both immunofluorescence and H&E staining were carried out as detailed in ^46^.

Antibodies are listed in Table S2. Immunofluorescence of Dystrophin was confirmed with antibodies from two sources (Table S2).

#### Cellular assays

Cellular assays were conducted in

*Fusion index*: cell cultures were vehicle-treated or Dox-induced cell cultures for 4 days, followed by immunofluorescence with the MF20 antibody, marking differentiated cells ^47^. Fusion index was calculated from the proportion of nuclei in MyHC objects. Fusion index is a calculated value which is measured in muscle cell culture cultivated under differentiation.

*Multi-nucleated cells*: were determined by the segmentation of nuclei in large objects containing >=3 nuclei. The remaining nuclei are mononucleated (Figure S2).

*Oligo-dT in situ hybridization to detect bulk poly(A) RNA*: cells were cultured in 96 Well Black/Clear Bottom Plate (#165305, ThermoFisher), and were fixed using 3.7% formaldehyde for 15 minutes at RT. After two PBS washes, the cells were incubated with protease III (#322337, Advanced Cell Diagnostics), diluted 1:30 in PBS, for 15 minutes at RT. After two washes with PBS, cells were incubated in a hybridization buffer (#10369, Cepham Life Sciences) for 15 minutes at RT. The 5’-Cy5-Oligo-dT12-18 probe (#26-4400-02, Gene Link) was diluted 1:1000 in hybridization buffer and incubated overnight at 40°C in a humidified chamber. The following day, washes were carried out at 40°C for 5 minutes in the following order: 4x, 2x, and 1x SSC buffer and with PBS. Finally, the cells were incubated with Hoechst and kept in PBS during imaging.

*Mitochondrial activity*: Tetramethylrhodamine, Methyl Ester, Perchlorate (TMRM) (#T668, ThermoFisher) staining was carried out in living cells according to the manufacturer protocol. Hoechst (#34580, ThermoFisher) nuclei counterstaining was added after TMRM incubation. Live cells were imaged at 560nm (TMRM) and 380nm for the nuclei.

### Cell growth and cell migration assays

Cell proliferation and migration were assessed using the IncuCyte® Live-Cell Analysis System (Sartorius). For proliferation, cells were seeded in a 96-well plate, and confluence was monitored over time using phase-contrast imaging with the standard IncuCyte® protocol. Values were normalized to time=0. For migration, at near-confluency (>90%), a scratch wound was introduced using the IncuCyte® WoundMaker™ tool, and wound closure was monitored by phase-contrast imaging at 3 hours intervals. Values were normalized to time=0. Images were acquired and analyzed using IncuCyte® software.

### Cell toxicity

Cell toxicity was assessed using the CellTox™ Green Cytotoxicity Assay (#TM375, Promega) according to the manufacturer’s protocol. CellTox™ Green dye was diluted in the appropriate growth medium and added directly to cells at the time of seeding. Fluorescence intensity, proportional to cell death, was measured over time using the IncuCyte® Live-Cell Analysis System (Sartorius). Values were normalized to time=0. Images were acquired and analyzed using IncuCyte® software.

### Imaging platforms and image quantification

The CellInsight CX7 LZR high-content screening (HCS) platform was used for high-content imaging. The accompanying HCS Platform Spot detector and Co-localization toolbox (ThermoFisher) performed a cell-based analysis.

Imaging was used to calculate the differentiation index, which was done with a 10x objective covering over 12,000 nuclei per well, using the Co-localization toolbox. Example of multinuclear segmentation in differentiated cell culture is in Fig. S10.

Imaging for PABPN1 quantification, oligo-dT, and TMRM was made with a 20x objective, covering at least 5000 nuclei per well. Puncta were calculated with the Spot detector toolbox. Nuclear FI was measured with the Co-localization toolbox. Quantifications of fluorescence signal in mouse cross-sections were carried out in ImageJ.

*PABPN1 aggregates* – first the nuclei were threshold from the DAPI channel, and the nuclei region of interest (ROI) were counted. PABPN1 puncta was masked with a stringent threshold, and the nuclear ROI was projected onto the PABPN1 mask (Fig. S11A). The puncta within nuclei ROI were counted, and their proportion was calculated from the total nuclei objects. *Dystrophin area* – equal regions, covering ∼1/3 of the cross section, were selected from each mouse. The dystrophin signal was threshold (Fig. S11B), and the proportion of the masked area was documented. An equal threshold was used across all samples. *MyHC-2A area and MFI* were measured from the entire muscle cross section. The cross-section area was measured from the Dystrophin signal with a low threshold, MyHC-2A in myofibers was masked, using a constant threshold across all samples (Fig. S11C). The proportion of the masked area and MFI were calculated using the cross-section area ROI. Automated *myofiber diameter* quantification was performed using MyofibrilJ (https://imagej.net/plugins/myofibrilj) after excluding technical artifacts, such as tissue folds and other damaged regions. For each mouse, the average value from two regions was used for analysis.

### Statistics and bioinformatics

Statistical tests for cellular and molecular assays (N=3 or 4 biological replicates) were made in GraphPad Prism 9.3.1.

Protein-protein interaction networks were identified with k-means clustering in STRING (https://string-db.org/) ^48^. Lollipop plots of the enriched clusters were made in STRING.

Venn Diagram was made with DeepVenn (https://www.deepvenn.com/) ^49^. Protein structure prediction was made with Alphafold 3 ^50^ using the entire protein.

### Data deposit

Mass spectrometry raw files have been deposited into the ProteomeXchange consortium via PRIDE with a reference number PDX0000 ^51^.

RNAseq raw files have been deposited into the European Genome-phenome Archive, reference nr (EGAX0000).

## Acknowledgments

We thank Leiden Genome Centre (LGTC) for the 1C library prep, and Leon H. Mei (Biostatistics) for assisting with RNAseq processing and APA-shift analysis. We also thank Jelle J. Goeman for his advice on biostatistical analysis. This study was partly funded by Human Genetics, LUMC.

