## Supplementary material for "Distinct roles of IDR and CCD domains control PABPN1 aggregation and enable therapeutic rescue": Suppl data

**Supplementary Materials**

**Table S1. List of primers**

|  | **Forward 5’** | **Reverse 5’** |
| --- | --- | --- |
| **Primers for generation of PABPN1 variants** | | |
| dAla-FLAG | TCGCAGTACTAAGATCTCCATCCAATGGGGGCTGCGGGCGGTCGGGGC | ATTGGATGGAGATCTTAGTACTGCGA  TGAATTCGAAGCTTGAGCTCG |
| dIDR-FLAG | TCGCAGTACTAAGATCTCCATCCA  ATGGAGGACCCGGAGCTGGAAGCTATC | CATTGGATGGAGATCTTAGTACTGCGA  TGAATTCGAAGCTTGAGCTCG |
| dCCD-FLAG | TCGCAGTACTAAGATCTCCATCCA  ATGCCTCCAGGCAATGCTGGCCCG | CATTGGATGGAGATCTTAGTACTGCGATGAATTCGAAGCTTGAGCTCG |
| trPAB-FLAG | AGCTCAAGCTTCGAATTCAATGGAGGAAGAAGCTGAG | TGAATTCGAAGCTTGAGCTCGAGATCTG |
| A16-FLAG | GACTACAAGGATGACGATGACAAGTAAATTAGGAGGAGAGAGAGG | TTACTTGTCATCGTCATCCTTGTAGTCGTAAGGGGAATACCATGATG |
| **RT-qPCR** | | |
| HPRT1 | TGGTCAGGCAGTATAATCCAAAGA | TCAAATCCAACAAAGTCTGGCTTA |
| FLAG | ACAGTGGTTTTAACAGCAGGC | CTTGTCATCGTCATCCTTGTAGTC |
| trPAB | GGAAGAAGCTGAGAAGCTAAA | CTTAACTCGCCATGGAATGA |
| PABPN1_3’-UTR | ATGGACACGTCTCAACTGCG | ACGGAGGGAAGGTAACAAGC |
| **Distal (D) and proximal (P) primer sets for APA analysis in cancer and muscle** | | |
| D_ASB5 | CACTATTTTCAACAACCCAGGGC | AGCTTAGTGATCTGGGGGTC |
| P_ASB5 | ACCCCAAGCTCTCTTTACCA | AGTAACGTTGGC AGCTGGAG |
| D_TMEM123 | CTTCGTAGACCGACCTGAAGT | TGCAGAGGAAACTTTCAAGGCA |
| P_TMEM123 | ACTCAAGAAGAGGCATTCGGT | TGATAGGGCAGCATCAATCTGT |
| D_CYBRD1 | AAAGCCTTTTCTCTCAAAGCG T | TGACACAAGCAGCACAGGAA |
| P_CYBRD1 | GGATGAGGCTGGGCAGAGAT | GGCTAATAGGAGAAGCAAAACTGT |
| D_LRP1 | AAGACGTGGCTCTGGGTGAG | CTCATAGGTGCCCCTGGAGTTG |
| P_LRP1 | GCACGGACGAGAAGCGAGAA | GCTTTCTGGGGGCAGTCC |
| D_PSTPIP2 | CACCTGTGCACTATGAGAAAATGG | CTGCACAGAGCAGGAGAGGAA |
| P_PSTPIP2 | TTTCCGGCTAGTGCTTCTGT | ATAGCAAGGGTTCACTTCAAAGTC T |
| D_CDKN1A distal | GTGAGGGTCCCATGTGGTG | GCCAGTGTCTCCCTCCTAGA |
| CDKN1A proximal | ACTCTCAGGGTCGAAAACGG | ATGTAGAGCGGGCCTTTGAG |
| **Distal (d) and proximal (p) primer sets for APA analysis in mice** | | |
| D_DMD | TGGCATGTATCATCATTGTCTCC | TGTTGTGAAAATGCAGTAAAACTGA |
| P_DMD | TGGCAGATGATTTGGGCAGA | CCATGCGGGAATCAGGAGTT |
| D_Oaz2 | CTGGACTGCAGAAATGGCCT | ACACCCCCACACCCAATGTA |
| P_Oaz2 | TACCCCCTGGACCAGAACTT | TCCCTAGACAGCCAG AGACC |
| D_Tmed3 | GGATCTTGCAAGGGGAAAACTAC | ACCAACAAACTGCTGCAAGG |
| P_Tmed3 | TTTGAAGTCCGGAGGGTTGT | TGACCAAGGTGTTTGGTAAACAAG |
| D_Phtf2 | AGTAAGTGACATCCCTGCCTT | CAGCACACAGCATTTGCCAT |
| P_Phtf2 | TTGTTGAAAGGGTGGCAGAGC | TGCTTAGCAATGGCTTCTGTT T |

**Table S2. List of antibodies**

| **Primary antibodies** | | | | | | | |
| --- | --- | --- | --- | --- | --- | --- | --- |
|  | **Dilution** | | | | **Host** | **Supplier** | **Cat. Number** |
|  | **Immunofluorescence (IF)** | | **Western blot (WB)** | |  |  |  |
| GAPDH | - | | 1:5000 | | Mouse | Invitrogen | MA5-15738 |
| PABPN1 (PAB2) | 1:1000 | | 1:4000 | | Rabbit | Homemade (10) |  |
| FLAG | 1:1000 | | 1:1000 | | Mouse | Sigma-Aldrich | F1804 |
| RPS3 | - | | 1:500 | | Mouse | Proteintech | 66046 |
| MATR3 | - | | 1:1000 | | Rabbit | Invitrogen | 68280 |
| MyHC | 1:250 | | 1:200 | | Mouse | DSHB | MF20 |
| MYOG | 1:1000 | | - | | Mouse | BD Pharmingen | 556358 |
| IGF2BP1/2 | - | | 1:200 | | Mouse | DSHB | PCRP-IGF2BP1-2D4 |
| Dystrophin | 1:1000 | | - | | Mouse | Gene Tex | GTX27163 |
| Dystrophin | 1:250 | | - | | Mouse | DSHB | 6A9 |
| **Secondary antibodies** | | | | | | | |
| **Application** | **Fluorophore** | **Dilution** | | **Supplier** | | | |
| IF | Anti-R-Cy5 | 1:2000 | | ThermoFisher Scientific | | | |
| IF | Anti-M-Alexa488 | 1:2000 | | ThermoFisher Scientific | | | |
| IF | Anti-M-Alexa594 | 1:2000 | | ThermoFisher Scientific | | | |
| WB | Anti-R/M-800/680 | 1:10000 | | IRDye 800CW or IRDye 680RD (LI-COR) | | | |

**Table S3: FLAG-trap protein binders in muscle and blader cancer cells**

**Table S4: Differentially expressed and APA-shift transcripts in trPAB bladder cancer cells**

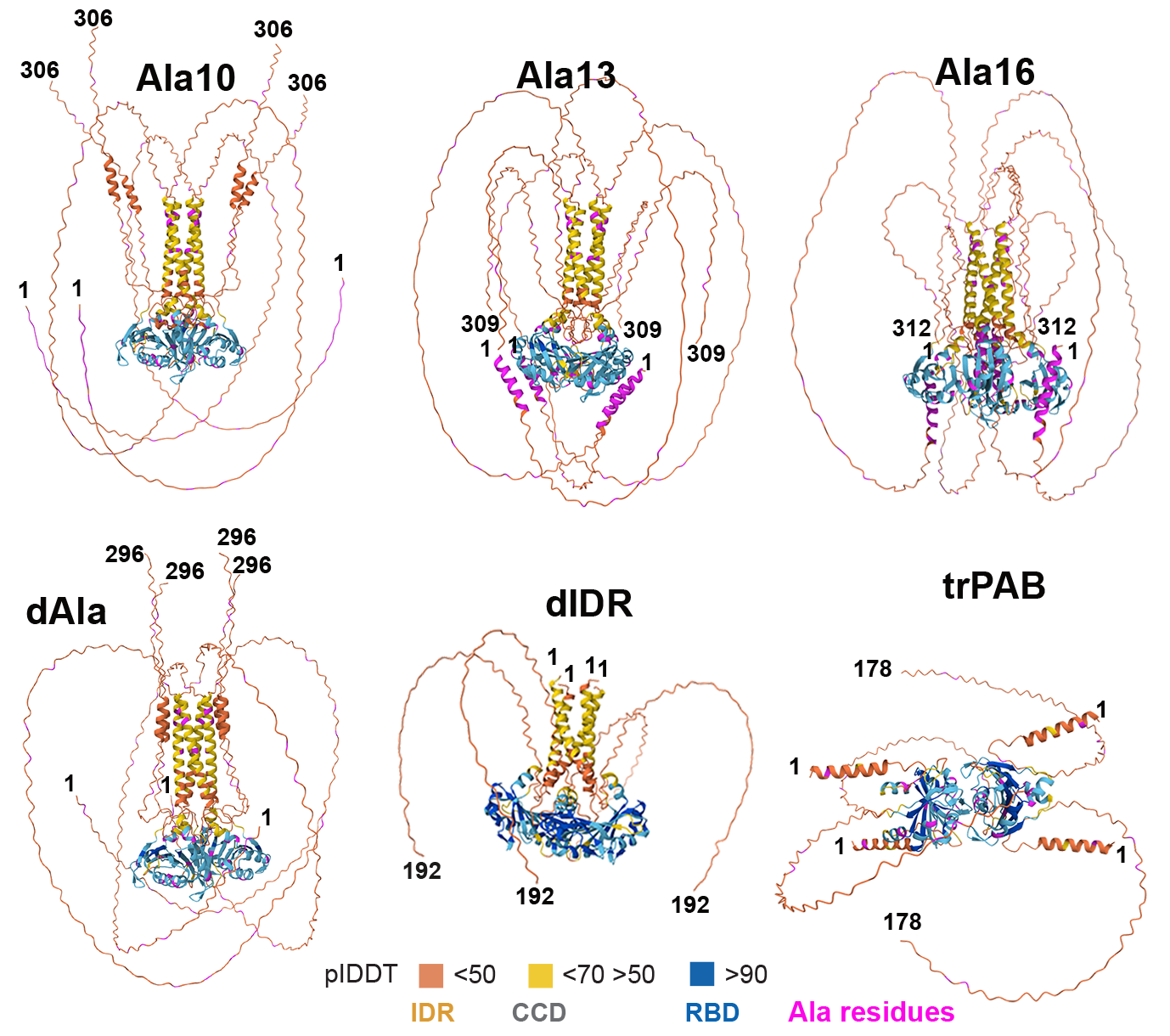

**Figure S1. AlphaFold prediction of full-length PABPN1 (upper row) and truncated isoforms (lower row).** The full amino acid sequence was used for all predictions, with the positions of the first and last residues indicated. Predicted local distance difference test (pLDDT) scores are shown as a colour scale: blue (high confidence, RNA-binding domain), cyan to yellow (moderate to low confidence, coiled-coil domain), and orange (low confidence, intrinsically disordered region). Alanine residues are highlighted in pink. In all models the orientation of the RBD is horizontal.

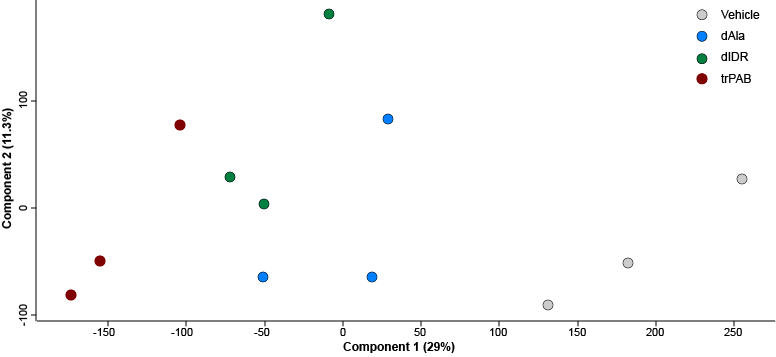

**Figure S2. A PCA plot of the 12 samples in muscle cells.**

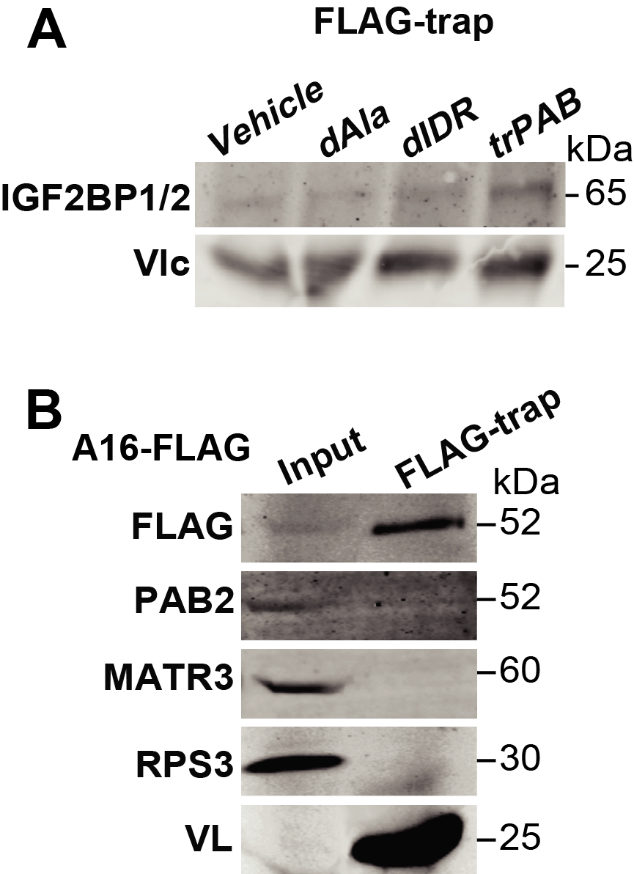

**Figure S3. Immunoblot of FLAG-trap.**

A. Immunoblot of FLAG-TRAP with anti-IGF2BP2 in muscle cells. **B.** Immunoblot of input and FLAG-trap in A16-FLAG cells with anti-Flag, PABs, anti-MATR3, anti-RPS3 antibodies. Molecular weight in kDa are depicted. The variable light chain (V) shows IP control.

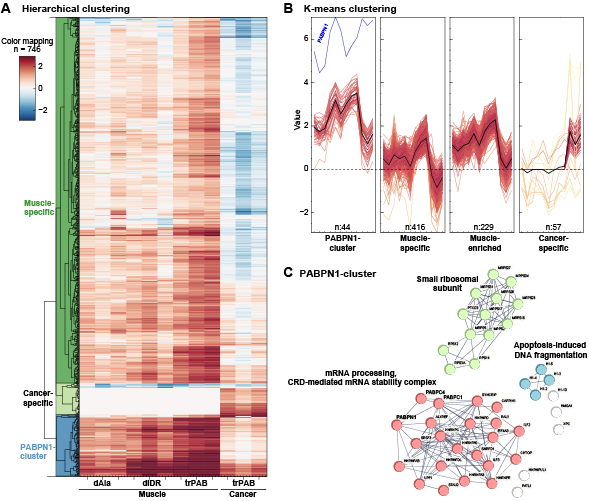

**Figure S4: Clustering of the PABPN1-interactome across different cell types**

Clustering was made with the fold-change (log2) values in muscle: dAla, dIDR, trPAB, and in cancer trPAB. **A.** Hierarchical clustering. **B.** K-means clustering. **C.** Protein-protein interaction network enrichment analysis of proteins within the k-means PABPN1 clusters. Clusters are color-coded, while non-clustered proteins are shown in white. PABPN1, PABPC4, and PABPC1 are highlighted.

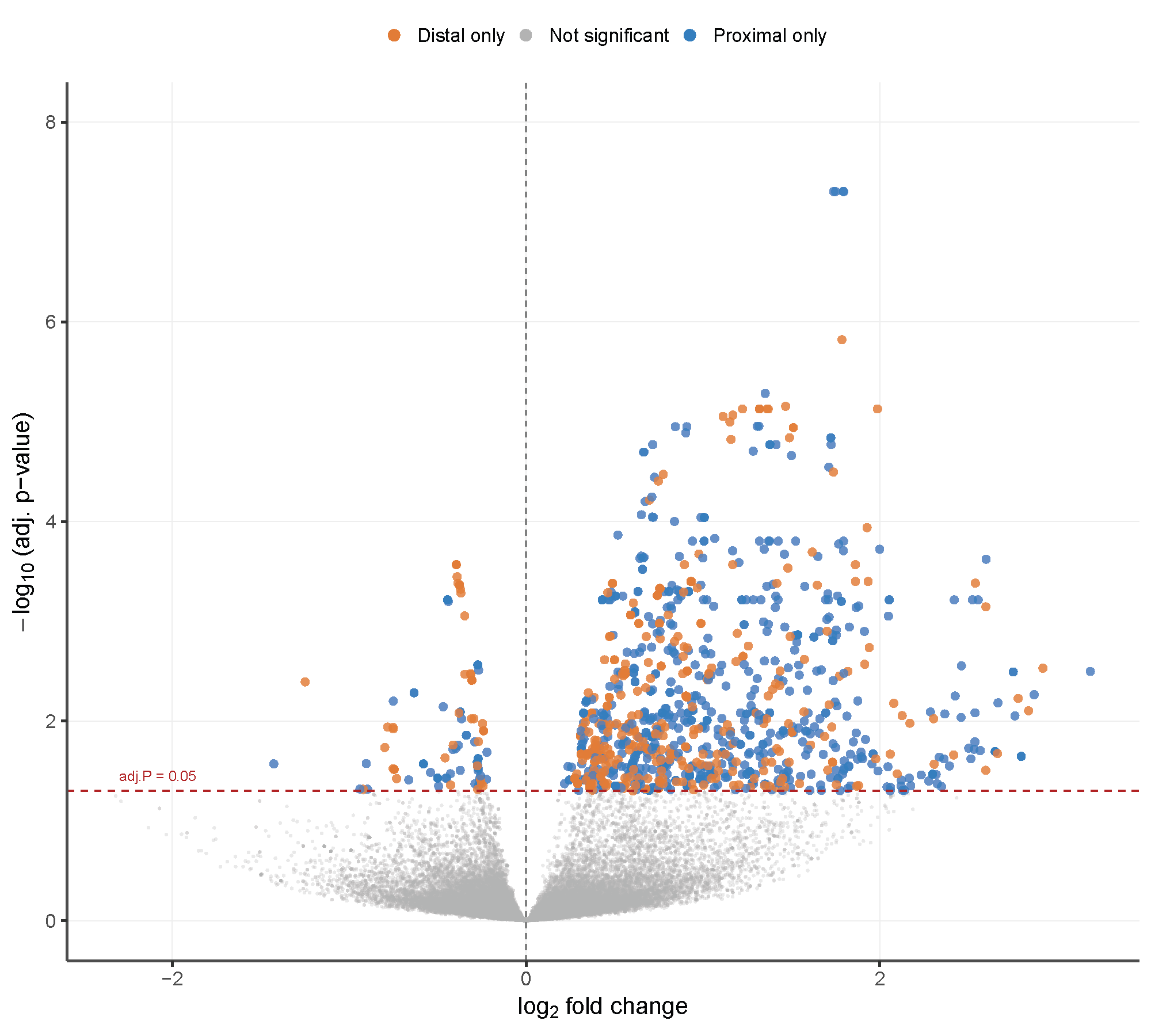
**Figure S5. Vehicle and trPAB in cancer cells 1C RNAseq.** Volcano of DE analysis at the 3’-UTR of transcripts tr-PAB vs. Vehicle. Positive fold change and colored shows significant higher levels in tr-PAB, P<0.05, FDR.

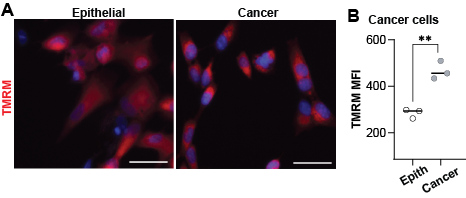
**Figure S6. Mitochondrial activity. A-B.** Epithelial and bladder cancer cells. **A.** Representative images are shown, with TMRM in red and the nuclear counterstain in blue. The Scale bar is 50 μm. **B.** The dot plot displays TMRM mean fluorescence intensity (MFI). Each dot represents a biological replicate, averaging ~3,000 cells. Statistical analysis was performed using one-way ANOVA; p < 0.01 (***).

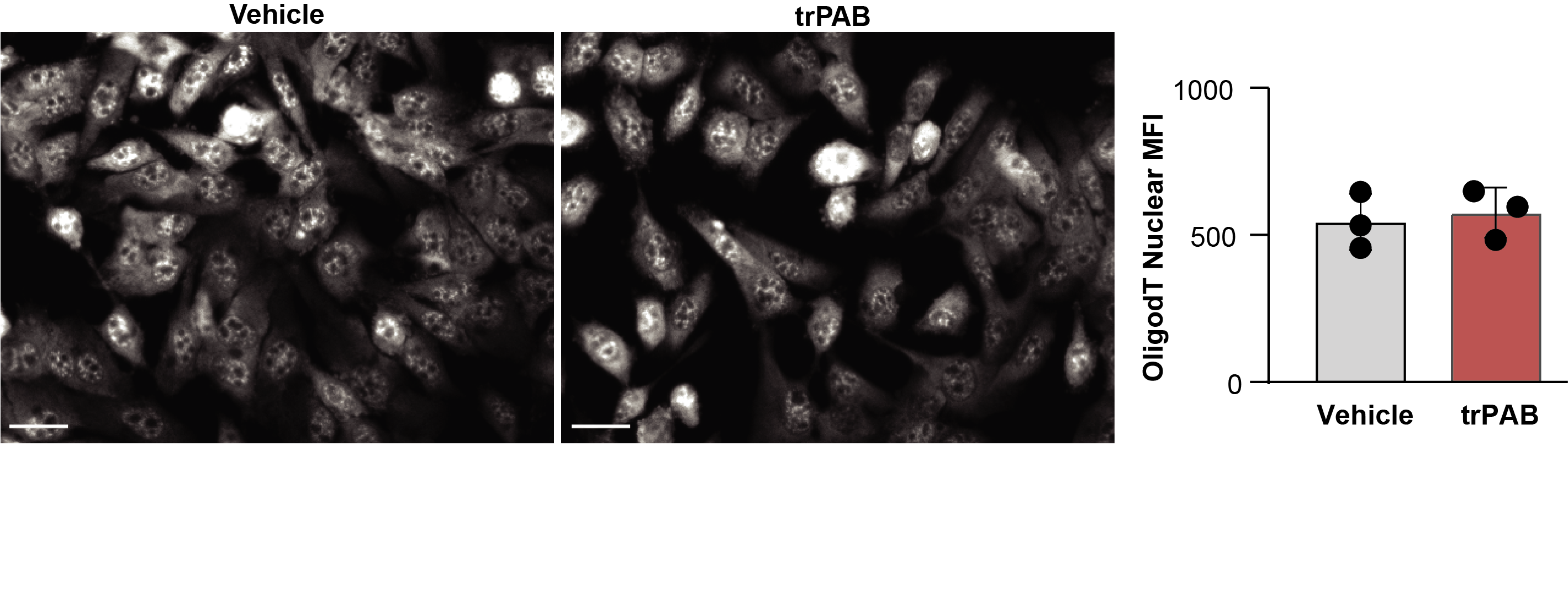
**Figure S7. Oligo-dT in situ hybridization in cancer expressing trPAB. A. Cancer cells.** Representative images show oligo(dT)-Cy5 staining in vehicle- and trPAB–treated cancer cells. Scale bar: 20 µm. Box plot shows oligo-dT MFI in the cell nucleus. Each dot represents a biological replicate of >1000 cells per replica.

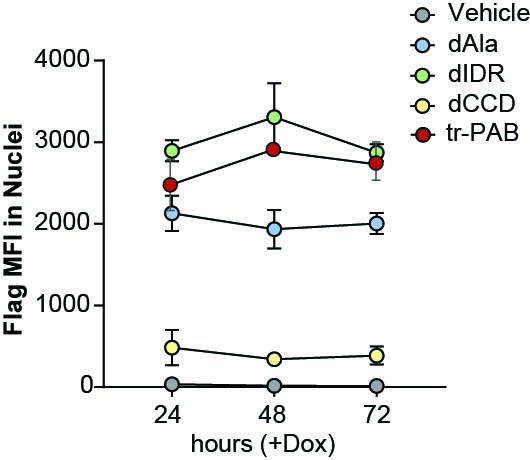

**Figure S8. Accumulation of Flag signal over time. Cancer cells were treated with Dox for the indicated time. Flag** signal was detected by immunofluorescence and was measured using the Cell-Insight imaging. Average and standard variation are from N=3 replicates.

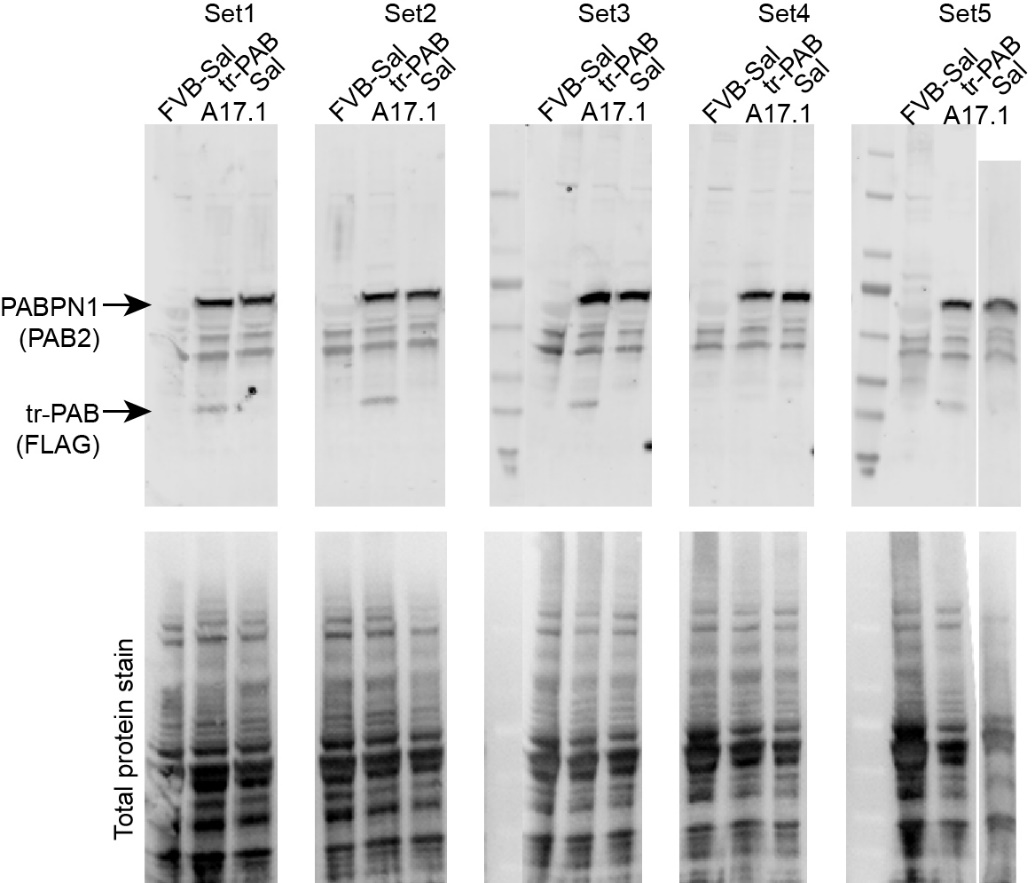

**Figure S9. Supplementary to Figure 5C: Western blots of protein extracts from TA muscle.** Blots were probed with PAB2 and anti-FLAG antibodies. A total protein stain is shown as the loading control. For each genotype, N = 5 mice were analyzed, loaded across five sets.

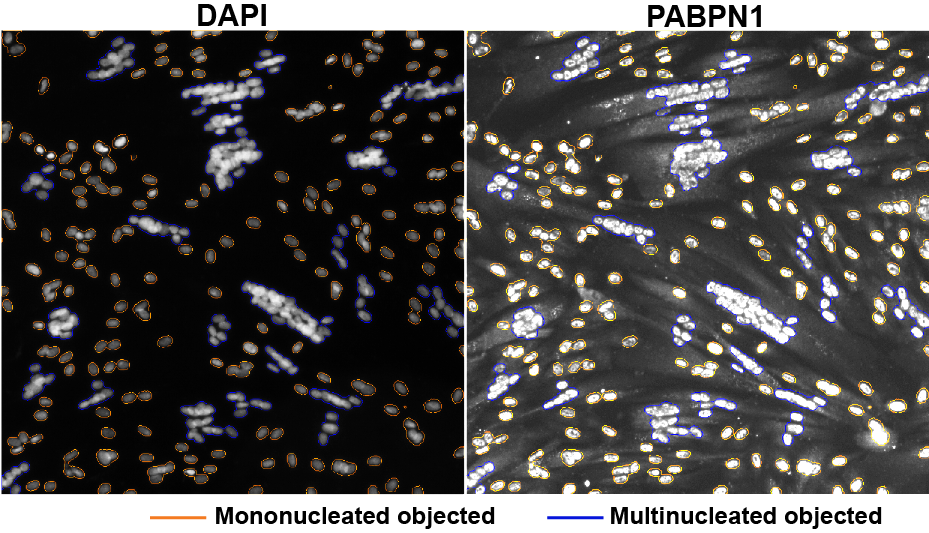

**Figure S10. An example of nuclear segmentation in differentiating cell cultures using nuclear counterstaining.** PABPN1 appears elongated and shows cytoplasmic localization. Multinucleated objects are outlined in blue, and mononucleated objects in orange.

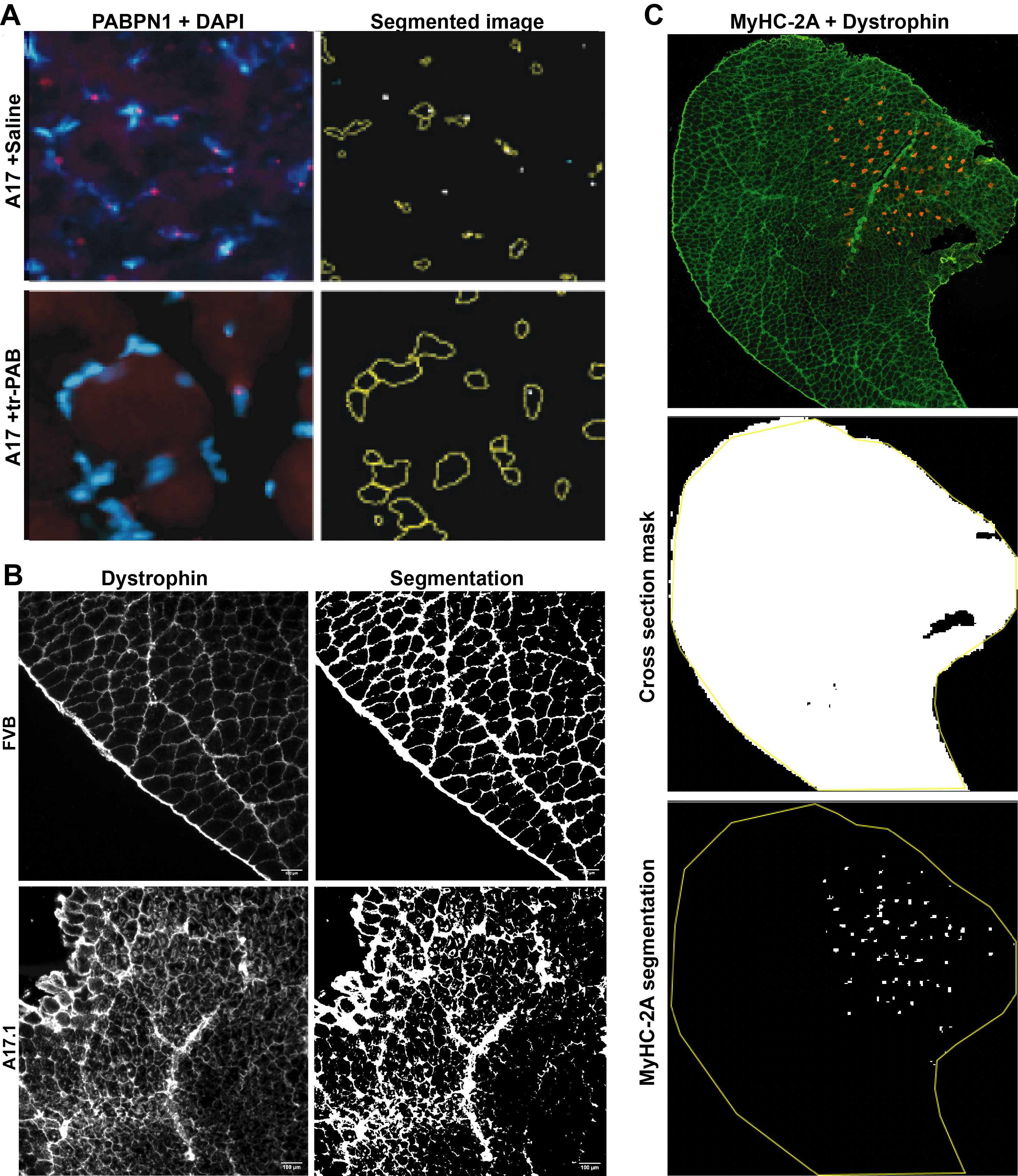

**Figure S11. Image analysis in muscle cross sections.**

**A.** Representative images (left) showing PABPN1 and DAPI signals, with corresponding segmentation (right) in A17 + saline– and A17 + tr-PAB–treated muscle. Nuclei areas are outlined in yellow, and PABPN1 puncta are indicated by white dots. **B.** Dystrophin, staining (left) and mask (right) in FVB and A17.1. **C.** MyHC-2A and dystrophin staining (top), cross-sectional area mask generated from the dystrophin channel (middle), and MyHC-2A segmentation with the cross-sectional area ROI outlined in yellow (bottom).
